# Strong mnemonic prediction errors increase cognitive control, attention, and arousal

**DOI:** 10.1101/2025.10.02.680173

**Authors:** Alice M. Xue, Jenna Jokhani, Anthony M. Norcia, Anthony D. Wagner

**Author notes:** **Corresponding author:** Alice M. Xue.

## Abstract

Ongoing experience is continuously processed in the context of past events. Divergence between current experience and memory-based predictions (i.e., a mnemonic prediction error; MPE) is theorized to be a signal that encoding, as opposed to retrieval, should be prioritized. We asked how MPEs place demands on control and attention-related processes, and whether these demands differ as a function of prediction strength. We investigated these questions across two experiments, wherein we recorded scalp electroencephalography and/or pupillometry as 101 young human adults performed an associative memory task. Strong MPEs, more so than weak MPEs, increased physiological indices of cognitive control (frontal theta), attention (posterior alpha and pupil size), and arousal (pupil size). Greater cognitive control (frontal theta) during strong MPEs promoted better learning of prediction violating (i.e., unexpected) stimuli. Collectively, these findings reveal multiple pathways through which the mind and brain respond adaptively to violations of strong mnemonic predictions.

## Introduction

Internal models of the world are constructed using memories of past experiences. Knowledge retained from events in the past enables predictions about ongoing and future events, allowing for adaptive behavior. As events unfold, alignment between what is experienced and what is predicted provides validation for the current internal model and signals that continuing to retrieve information from memory may be useful for guiding behavior. On the other hand, divergence of ongoing experience from mnemonic predictions signals that the environment has potentially changed and that past experiences may not be informative or relevant for current and future behavior. Such mnemonic prediction errors (MPEs) may elicit increased attention and arousal^1^ and signal that more cognitive control –– the collection of cognitive and neural mechanisms that effortfully influence information processing to be aligned with one’s goals^2^ – is needed.

Insights into the effects of MPEs on attention and arousal have come from studies of event segmentation. According to Event Segmentation Theory^3,4^, the mismatch between one’s internal model and ongoing events can elicit a prediction error, leading to event segmentation (i.e., the psychological perception of an event boundary along with accompanying neural computations), which discretizes current experience into “episodes” in memory^5,6^. Extant data indicate that increases in pupil size are observed at event boundaries^7,8^ and in response to prediction errors of varying content, signs, and modalities^9–11^. Such changes in pupil size reflect increases in attention and arousal^12–14^ and may be linked to noradrenergic activity in the locus coeruleus (LC)^14,15^. The LC has been associated with signaling strong prediction errors^16,17^, also characterized as “global model failure signals”^16^ that initiate a network reset^18^. The LC may further support adaptive behavior in the face of MPEs by enhancing memory encoding to facilitate model updating^19^. New learning following MPEs may depend on increases in attention, by direct modulation of hippocampal learning mechanisms^19,20^, or both. While MPE responses in the hippocampus appear to vary with prediction strength^21^, it is unclear whether effects on attention and arousal are similarly modulated. Here, we hypothesize that pupil size will show a greater increase in response to strong compared to weak MPEs, that MPE-induced increases in attention and arousal will support learning, and that learning from MPEs will differ as a function of prediction strength^11^.

Complementing evidence of MPE-related increases in pupil size^7,8,11^, time-resolved electrophysiological measures of visuospatial attention –– specifically, electroencephalography (EEG) posterior alpha power suppression^22,23^ –– may also show that attention increases following MPEs, evidenced by greater alpha suppression^24^. The underlying sources of these measures of attention may be partially shared, given that pupil size and posterior alpha are coupled in certain contexts^25–27^. Moreover, LC lesions in monkeys reduce the amplitude of the P3 event-related potential (ERP), another electrophysiological measure of attention/alertness^28^. However, it remains unclear whether LC activity additionally modulates posterior alpha suppression. Posterior alpha and pupil size also could be differentially influenced by other mechanisms of MPE detection, such as prefrontal cortex (PFC)^29^.

PFC theta, observed using magentoencephalography (MEG)^30,31^ and scalp EEG^32–34^, is known canonically as frontal theta^35^ or frontal midline theta^36^ and is considered a marker of cognitive control^35,37^. Frontal theta increases following reward prediction errors^32,34,35^, scales with negative prediction errors^32^, and fosters behavioral adaptation^32^. Behavioral adaptation in response to increases in frontal theta may be mediated by changes in information processing^35^ by way of increased attention. Control may directly increase attention to and prioritization of bottom-up sensory information from the external environment and the overcoming of memory-guided, but now irrelevant, representations and behavioral responses^38^. In rodents, PFC activity can modulate LC activity^39^, raising the possibility that the behavioral adaptation effects associated with increased frontal theta could be mediated in part by changes in LC activity and thus pupil-linked attention and arousal. Whether MPEs elicit (a) an increase in cognitive control that (b) consequently results in increases in attention and arousal to facilitate adaptive behavior remain open questions. A primary aim of the current study was to test whether there are interactions among cognitive control, attention, and arousal upon detection of MPEs.

Characterization of potential process interactions may additionally provide critical insights into mechanisms by which MPEs facilitate learning in service of future behavior. Higher learning rates following MPEs may be mediated, in part, by increases in attention and arousal^1^, which facilitate memory encoding^11,40,41^. In addition, increases in pupil-linked attention and arousal following MPEs could reflect direct involvement of LC activity in fostering learning, given that optogenetic activation of LC neurons in mice enhances memory encoding^42^ and higher LC integrity among older adults is associated with better episodic memory performance^43^. LC effects on learning may also be mediated by the hippocampus, which receives profuse projections from LC neurons^42^. Increases in attention and arousal following MPEs may require the initial detection that cognitive control, as instantiated via frontal theta^35^, is needed. Higher frontal theta prior to^44^ and during^37,45^ encoding has independently been associated with better subsequent memory, leaving open the additional possibility that MPE-related learning effects stem from changes in frontal theta. Finally, MPE-driven increases in frontal theta may also enhance learning by driving hippocampal theta^30^, which also is associated with better subsequent memory^46^.

There are thus multiple possible pathways by which MPEs impact cognition. Here we focus on two mechanisms: (1) increases in cognitive control (assayed via frontal theta) and (2) increases in attention and arousal (assayed via posterior alpha and pupil size). To elicit MPEs, we used an associative memory task wherein participants were tested on their memory for verb-picture pairings they had studied four times (strong memories) or once (weak memories). Associative memory of a paired picture was cued by the presentation of a studied verb. On a subset of trials, verb presentation was followed by the presentation of a mismatched picture. Onset of a mismatch would indicate a violation of a mnemonic prediction and the elicitation of an MPE^47^. Using time-resolved scalp EEG (Experiment 1) and pupillometry (Experiments 1 and 2) measured as participants completed the associative memory task, we document the effects of strong vs. weak MPEs on cognitive control, attention, and arousal. Then, we characterize how control signals interact with attention and arousal signals, and how EEG and pupil assays of attention/arousal relate to each other. Throughout, we examine how these electrophysiological and pupillary measures differ as a function of memory strength to investigate how trial-level assays of prediction strength relate to MPE responses. Finally, we examine whether MPE-related changes in each of these physiological assays relate to the encoding of prediction-violating stimuli, as evidenced by subsequent memory for mismatch probes.

Collectively, our findings (**Fig. 1**) show that strong MPEs, relative to weak MPEs, evoke greater increases in cognitive control (frontal theta), attention (posterior alpha suppression and pupil), and arousal (pupil). Exploratory mediation analyses did not provide strong support for the hypothesis that increases in attention and/or arousal induced by MPEs can be explained in part by increases in cognitive control. Finally, increases in frontal theta, but not posterior alpha or pupil size, are linked to enhanced learning of strong prediction-violating (i.e., mismatch) stimuli. As such, mnemonic predictions trigger increases in multiple neurocognitive modulatory processes, with error-driven increases in cognitive control fostering new learning.

**Figure 1.**
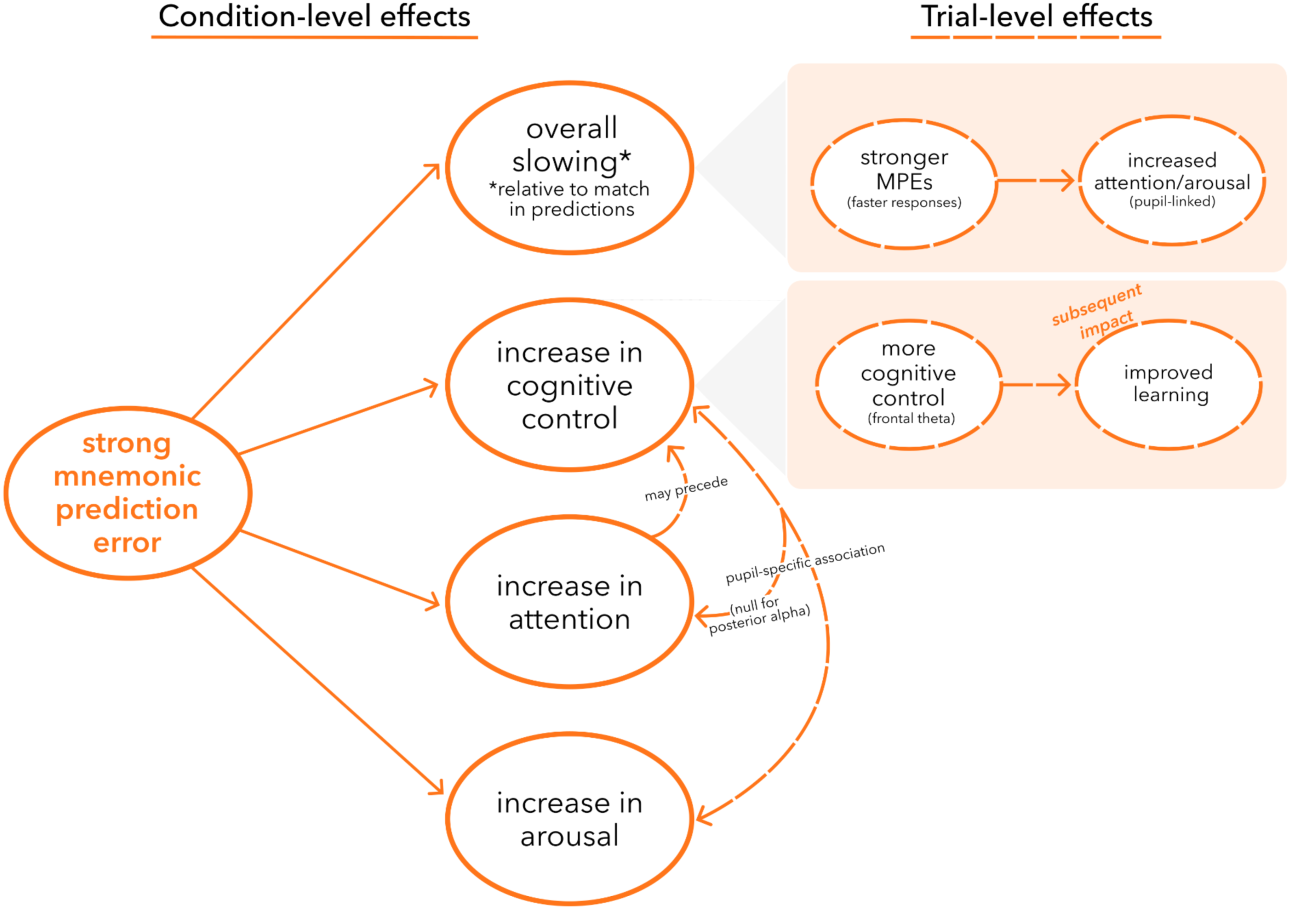
Summary diagram of the observed effects of strong MPEs in contrast to weak MPEs (solid lines in orange) and their effects at the trial level (dashed lines in orange).

## Methods

We conducted and, for most analyses, pooled data across two experiments that used a common associative memory task paradigm with pupillometry. Experiments 1 and 2 differed in: participant recruitment pool; the duration of the delay between the Associative Memory Task and a final subsequent Recognition Test on the mismatch probes; day of administration of the gradual continuous performance task (gradCPT); inclusion of a Localizer Task (only administered in Experiment 1); and concurrent recording of EEG (only collected in Experiment 1). Images of animate and inanimate objects were taken from the BOSS database^48^, an Animacy x Size stimulus set^49^, and the internet. Objects were isolated from their white or complex backgrounds and were then overlaid on a gray background for stimulus presentation. Scene stimuli in the Localizer Task in Experiment 1 were taken from the “Massive Memory” Scene Categories stimulus set^50^. Luminance of image stimuli in all tasks, with the exception of the gradCPT, were controlled using the SHINE Toolbox^51^.

### Experiment 1: Associative Memory Task with Same-Day Mismatch Probe Recognition Test (scalp EEG and pupillometry)

Fifty-one healthy young adults (aged 18-35 yrs) enrolled in Experiment 1. One participant withdrew early and all data from this participant were excluded. The final sample included 50 participants (25 female; mean age = 23.1 yrs, s.d. = 3.3). Pupil data were not properly tracked from three participants and were excluded from the corresponding pupil analyses. Participants were recruited from Stanford University and the surrounding community, were right-handed, and had normal or corrected-to-normal vision. The study was approved by the Stanford University Institutional Review Board and all participants provided written informed consent. In a single experimental session, participants completed 2.5hrs of behavioral tasks, with the subsequent recognition memory task occurring 6-min following the associative memory task. Throughout, EEG data were recorded using a 128-channel HydroCell Sensor Net (Electrical Geodesics) and pupillometry data were collected using an Eyelink 1000 eye-tracker; EEG and pupillometry data were collected at a sampling rate of 1000 Hz. Following EEG net application, electrode positions were recorded using a LiDAR sensor on an iPad using the Polycam application (https://poly.cam/). Before each task (ranging from 3–16min in duration), EEG impedances were checked and calibration and validation of the eye-tracker were performed.

### Experiment 2: Associative Memory Task with Delayed Mismatch Probe Recognition Test (pupillometry)

Fifty-three healthy young adults (aged 18-23 yrs) enrolled in Experiment 2. Participants were recruited from the Stanford Psychology Credit Pool and had normal or corrected-to-normal vision. Two participants withdrew on Day 1 and all data from these participants were excluded. One participant withdrew before Day 2 of the experiment; Day 1 data from this participant were retained. The final sample included 51 participants (23 female; mean age = 19.8 yrs, s.d. = 1.4). Other data loss/inclusion: one participant’s pupil was not properly tracked throughout the experiment and these pupil data were excluded from corresponding analyses; due to experimenter error, the subsequent memory test was administered incorrectly for four participants on Day 2 and these data were excluded from the corresponding analyses; one participant’s subsequent recognition data were excluded because the participant responded “Old” on all trials. The study was approved by the Stanford University Institutional Review Board and all participants provided written informed consent. In Session 1, participants completed 2hrs of behavioral tasks. Session 2 took place 48hrs after Session 1, during which participants completed the subsequent recognition task and the gradCPT. In both sessions of the experiment, pupillometry data were collected during behavioral tasks using an Eyelink 1000 eye-tracker at a sampling rate of 1000Hz. Calibration and validation were performed before each task.

### Experimental Design

#### 1.1 Associative Memory Task

Participants performed an associative memory task, which consisted of an associative encoding phase and an associative retrieval test. The associative memory task was administered in four study-test cycles, with encoding and retrieval phases separated by a 3min distractor task wherein participants reported the parity of sequentially presented integers ranging from 1 to 100. Each integer was presented immediately following a response. The purpose of the distractor task was to prevent participants from maintaining the studied pairs in working memory prior to the associative retrieval test.

In the associative encoding phase, participants were instructed to learn verb-picture pairs by forming a mental image involving the verb and the image in each pair (**Fig. 2a**). On each trial, an action verb was presented at the center of the screen for 1.5s. The subsequent inter-stimulus interval (ISI) ranged from 0.5-1.5s after which a picture of an animate or inanimate object was then displayed for 2.5s. During the presentation of the object, participants generated a mental image and reported (using a button press) whether their mental image of the verb-picture pairing was “very vivid” or “not very vivid.” The inter-trial interval (ITI) ranged from 1.25–1.5s.

**Figure 2:**
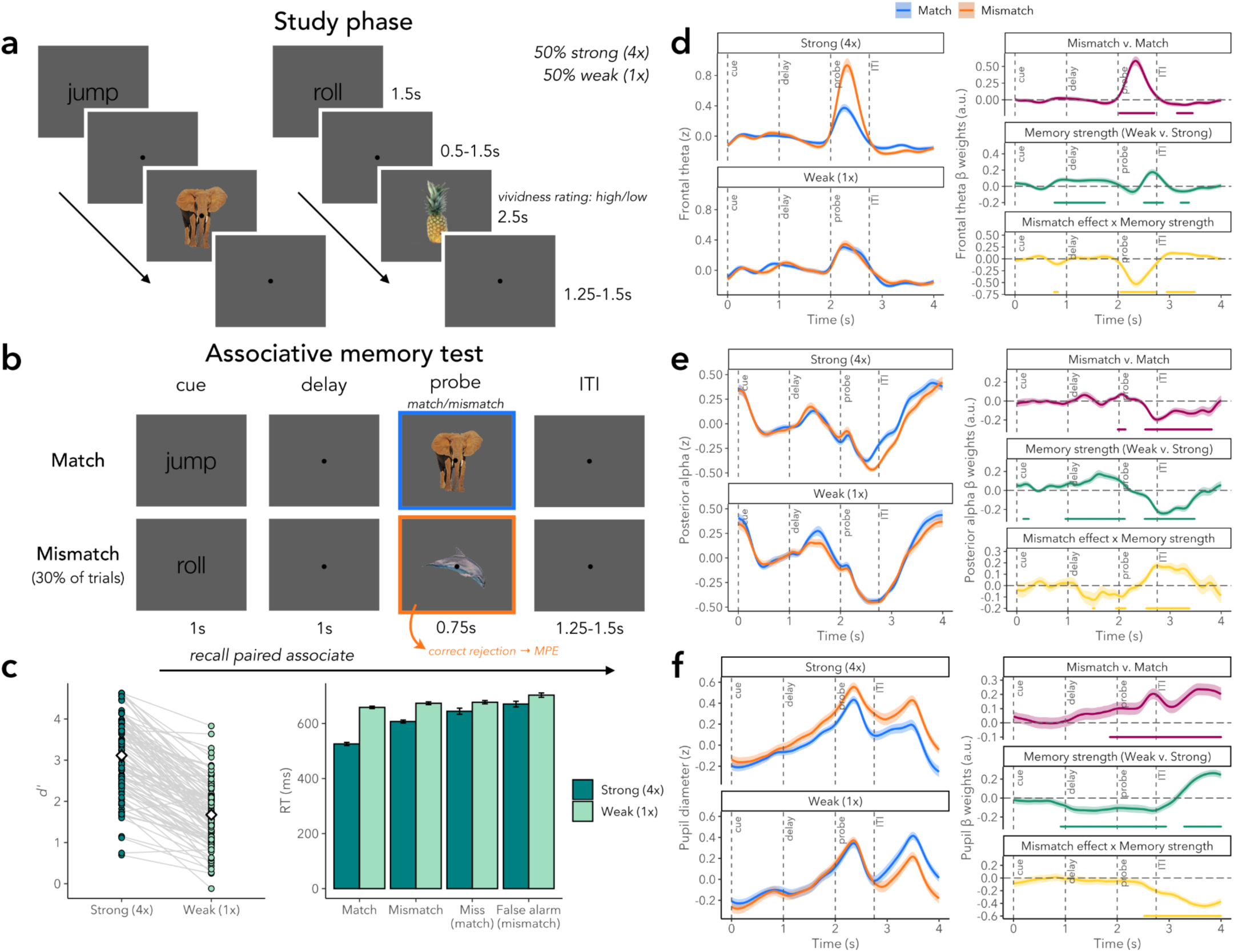
Associative memory task design and memory behavior. (a) Study phase task design. (b) Match/mismatch test phase task design. (c) Behavior in the match/mismatch retrieval task: *d’* (left) and RTs (right) in dark teal for strong pairings and light teal for weak pairings. Frontal theta power (d), posterior alpha power (e), and pupil diameter (f) for strong (top) and weak (bottom) trials, color coded by match/mismatch condition (blue/orange). Subplots on the left depict mean and standard error of z-scored power/size across participants. Mean and standard error of participant-level betas for the effects of Mismatch, Strength, and Mismatch x Strength are depicted on the right. Horizontal lines below beta plots indicate significant clusters (permutation-corrected; all p<0.001).

Participants were shown 52 pairs in each encoding phase; 26 were presented four times over the course of an encoding phase cycle (“Strong” pairings) and 26 were presented once (“Weak” pairings). Repeated exposures to strong pairings were distributed across the encoding phase cycle, such that each strong pairing was viewed once during each quartile of trials. As such, each cycle of encoding included 130 trials (26 pairs presented four times and 26 pairs presented once). A black fixation dot was presented at the center of the screen during all trials and participants were instructed to fixate throughout. Verb-picture pairings were randomly assigned per participant, as was the assignment of pairs to the Strong and Weak conditions.

After the distractor phase, an associative retrieval test examined memory for the verb-picture pairings (**Fig. 2b**). On each trial, a verb cue was presented for 1s. Participants were instructed to try to bring to mind the associated picture as soon as the cue appeared on the screen. Following a 1s delay, during which only the central fixation dot was presented on the screen, a picture was then presented for 0.75s. Participants were instructed to respond as quickly as possible, using a button press to indicate whether or not they thought they had studied the picture with that verb. Memory for all 52 pairs in the encoding phase of a given cycle was tested. The brief (0.75s) presentation duration of the test probe and instruction to respond rapidly were designed to encourage participants to retrieve the associate upon presentation of the verb cue, setting up a situation in which memory of the associate and perception of the probe either match or mismatch^47^. To focus on trials on which retrieval likely preceded probe onset, retrieval task trials were retained for analysis if responses were made within 1s after probe onset^47^. In each study-test cycle, 36 trials were “Match” trials, wherein the picture after the verb matched that studied in the encoding phase, and 16 were “Mismatch” trials, wherein the picture did not match what was studied. On Mismatch trials, the probe picture was from a different category as the correct associate (i.e., if the picture associated with a verb cue was animate, the mismatch probe was inanimate, and vice versa). To control for stimulus novelty during retrieval, mismatch stimuli were initially introduced to participants at the outset of the experiment in a pre-exposure 1-back task (see below). ITIs ranged from 1.25–1.5s. A black fixation dot was presented at the center of the screen at all times and participants were instructed to fixate throughout. The test order of pairings was randomized.

Each study-test cycle included unique verbs and pictures. The assignment of each picture to each verb for Match and Mismatch pairings was randomized across participants.

#### 1.2 Pre-Exposure Task

At the outset of the experiment, prior to the associative memory task, participants performed a 1-back task in which they were shown a sequence of images (each presented for 2s) separated by variable-length ITIs (1.0-1.2s). For each stimulus, participants were instructed to report, using a button press, whether the current stimulus was the same as or different from the previous stimulus. To maximize the duration with which participants attended to each stimulus, they were instructed to provide their response when each stimulus had left the screen. There were 96 unique stimuli in total (48 animate and 48 inanimate objects); 64 of the stimuli were later presented as mismatch probes during the retrieval phase of the associative memory task (see above), and all stimuli were later presented in the subsequent recognition test; the order of the stimuli was randomized. Of the 32 items not presented in the retrieval phase of the associative memory task, nine were repeated in the 1-back task. A black fixation dot was presented at the center of the screen at all times during all trials and participants were instructed to fixate on the central dot throughout the task. The purpose of this pre-exposure task was to familiarize participants to the stimuli that would later serve as mismatch probes during the associative memory task. By using mismatch probes that were familiar rather than novel, item novelty could not be a primary source of memory evidence guiding decisions in the retrieval phase of the associative memory task.

#### 1.3 GradCPT

To obtain a subject-level assay of attention, we collected 5-min of gradCPT^52^ data in each experiment with modified stimulus presentation timings. In Experiment 1, participants performed the gradCPT after four cycles of the associative memory task to introduce a delay before the surprise recognition test (see below) and prevent maintenance of material studied in the associative memory task in working memory. In Experiment 2, where the surprise recognition test took place 48hrs after the associative memory test, the gradCPT was performed after the recognition test in Session 2. In the gradCPT, a centrally presented picture gradually transitioned between different mountain and city images (90% of images were of cities and 10% were of mountains). Each image was presented for 0.05s and transitioned to the next image over 0.8s. Participants were instructed to press the spacebar when they saw a city scene and to withhold responding for mountain scenes. Performance in the gradCPT was not central to the core hypotheses and is not reported in the current manuscript.

#### 1.4 Recognition Test

To assess the effects of MPEs on learning, a recognition test was administered. In this surprise recognition memory test, participants were tested on their memory for the 64 mismatch probes shown during the associative retrieval test. Non-mismatch foils on the recognition test consisted of the 32 stimuli from the pre-exposure 1-back that were not repeated in that task nor used as mismatch probes during the associative retrieval test; as such, they were old but not associated with MPEs. An additional 64 novel foils were included in the recognition test. On each recognition trial, participants reported whether they had seen the image at any point during the experiment, by pressing one button for “old” and another for “new”. Each image was displayed for a minimum of 1.2s and a maximum of 2s, such that if participants failed to respond within 1.2s, the image remained on the screen for an additional 0.8s. ITIs varied from 0.5–1.5s and the order of the stimuli was randomized. A black fixation dot was presented at the center of the screen at all times and participants were instructed to fixate. In Experiment 1, the recognition test was administered after the gradCPT. In Experiment 2, the recognition test was the first task administered in Session 2. The recognition test examined subsequent memory for mismatch probes either ∼5-10min (Expt 1) or ∼48hrs (Expt 2) after the associative memory task.

#### 1.5 Localizer Task

At the end of Experiment 1, participants performed a second 1-back task designed to enable training of an EEG classifier that discriminates between evoked electrophysiological activity patterns associated with animate and inanimate objects. The localizer task was included in Experiment 1 to address other research questions and results from this task are not reported in the current manuscript. For completeness, the task is described here. On each trial, an image of an animate object, an inanimate object, or a scene^50^ was displayed on the screen for 2s. Images were separated by a 1.0-1.2s ITI. The instructions were the same as for the pre-exposure 1-back task, such that participants were instructed to report whether a given image was the same as or different from the previous image, and to respond after the image had left the screen. The stimuli were novel (i.e., had not appeared in any other experimental phase), with 270 unique stimuli presented in total (90 per category); the order of the stimuli was randomized. Ten percent of the stimuli repeated once (as 1-backs) over the course of the task.

### EEG and Pupillometry Analyses

#### 2.1 EEG Preprocessing

EEG data from the associative retrieval test in Experiment 1 are a focus of the present manuscript, given the open questions/hypotheses regarding mnemonic prediction errors. Preprocessing of these EEG data was performed using the MNE package^53^. EEG data were first downsampled from 1000 Hz to 250 Hz and then filtered with a high-pass filter of 1 Hz and a low-pass filter of 30 Hz. Bad channels were then identified via visual inspection of the power spectrum and were subsequently interpolated. An ICA was fitted to the data; ICs resembling blinks and muscle artifacts were identified automatically using functions in MNE and by visual inspection. Data were then re-referenced to the average.

#### 2.2 EEG Spectral Analyses

For analyses of spectral power in the associative retrieval task, trial-level power was computed on epochs of data from −1s to 5s with respect to trial onset. Epoched data were decimated by a factor of two to reduce computation time, yielding a sampling frequency of 125Hz. Epochs wherein the voltage of any channel of interest exceeded 100 μV were excluded from analyses. Frontal theta (4-7Hz) power was computed using electrodes 5, 6, 7, 11, 12, 13, 106, 112 and posterior alpha (8-12Hz) power was computed using electrodes 62, 67, 71, 72, 75, 76, 77 (**Supplementary Fig. 1**). These electrodes were selected based on prior research on attention lapsing and goal coding^54^. Instantaneous power was computed at each time point using a Morlet wavelet with 7 cycles and epochs were subsequently trimmed to 0s–4s after trial onset for subsequent analyses.

#### 2.3 Pupillometry Preprocessing

Pupillometry size data from both experiments were preprocessed^55^, with data first being decimated from 1000 Hz to 100 Hz. A pupil dilation speed threshold was determined by first computing the median absolute deviation (MAD) in pupil size in each block of associative retrieval task data. Visual inspection was used to assign 10 as the constant value for multiplicatively scaling the MAD. A rejection threshold was then determined as the sum of this value and the median dilation speed. Samples of data for which the change in pupil diameter from the previous sample exceeded this threshold were removed. Blinks were padded by 50ms on each side and linear interpolation was used to fill in missing data due to blinks or dilation artifacts. The data were then smoothed using a third order Butterworth filter and additional outlier samples were identified using the dilation speed threshold procedure described above, but with a constant factor of 0.2. This procedure of interpolation, smoothing, and outlier detection was then performed a second time. Finally, trials were excluded if more than 10% of data points were missing in the original time series. Data were subsequently de-trended by taking the residuals of a linear regression model fit to the time series for each run of each task. For all analyses, the z-scored residuals were analyzed.

#### 2.4 Statistical Models for Time Series Analysis

Trial-level regression analyses were conducted on the frontal theta, posterior alpha, and pupil time series from the associative retrieval test to identify temporal clusters that were sensitive to the factors of Mismatch (i.e., mismatch vs. match probes) and/or Strength (i.e., strong vs. weak associative pairs). For each participant and each time point, a linear regression model testing main effects of Mismatch and Strength, and a Mismatch × Strength interaction was run using R^56^ to compute beta weights for each regressor. For these models, only correct trials (i.e., hits to match probes and correct rejections to mismatch probes) were included to identify temporal clusters associated with memory retrieval strength and/or MPEs. For frontal theta and posterior alpha analyses, two participants in Experiment 1 were excluded because they were missing trials in at least one of the four conditions and model fitting could not be performed; for pupil analyses, two participants from Experiment 2 were excluded for missing trials in at least one of the four conditions. For analyses of subsequent recognition memory for mismatch probes, 10 participants from Experiment 1 and seven from Experiment 2 were excluded for missing trials in at least one of the four conditions; of these, seven participants from Experiment 1 were excluded from the frontal theta and posterior alpha time series analyses of subsequent recognition memory. For primary pupil analyses, data were pooled across Experiments 1 and 2 (N=94); see Supplement for results by experiment.

Significance testing for each regressor was conducted at each time point and was performed using one-sample t-tests. The beta weights computed for each time point were averaged across participants for visualization of the effects of Strength, Mismatch, and Mismatch × Strength interactions for frontal theta, posterior alpha, and pupil size over time. Contiguous time points with significant t-values were identified; the significance of these temporal clusters was then tested using permutation tests in which the sign (positive or negative) of each participant’s beta weights was randomly assigned; this procedure was repeated 1000 times and t-tests for each time point were computed on each iteration to generate a null distribution. Significance testing for each temporal cluster was performed by computing the probability of a cluster that was at least as long in the null distribution.

#### 2.5 Pupillary Temporal PCA

In addition to canonical pupil size analyses, the pupil data were deconstructed using a temporal principal components analysis (PCA). Temporal PCA was used to obtain trial-level assays of attention and arousal (1) with greater sensitivity to cognitive processes engaged during the task and (2) that could be related to electrophysiological measures on a trial-by-trial basis. PCA was performed on the pupil time series aggregated across all correct trials and all participants (N=96); the size of the input matrix was the number of trials × the number of time points (10346 × 401 time points). We retained the first six components, using at least 95% variance explained as the cutoff; cumulative variance explained by all six components was 96.16%. To obtain interpretable scores for each trial and each PC, the loadings were then orthogonalized using a varimax rotation on the rotation matrix produced by the PCA. For each component, we fit linear mixed effects models using lme4^57^ to examine the effects of Mismatch and Strength (score ∼ Mismatch * Strength + (Mismatch * Strength | participant)). We also examined how scores related to reaction time (RT) on hit and correct rejection trials (RT ∼ score ∼ Mismatch * Strength + (Mismatch * Strength | participant)). RTs were log transformed in all analyses to account for skewed distributions. A maximal random effects approach was adopted as recommended by Barr^58^; for models that failed to converge, we successively simplified the random effects structure.

#### 2.6 Multiple Mediation Analyses

To characterize the dynamics between cognitive control, attention, and arousal following a strong MPE, Bayesian multivariate mediation analyses^59^ were conducted using the brms^60^ package in R. All models were conducted on strong Mismatch trials. The first model tested whether the relationship between MPE magnitude, approximated using strong mismatch RTs, and immediate pupil MPE effects (PC3) was mediated by frontal theta effects (mean power in the second Mismatch × Strength cluster in **Fig. 2d**). The second model tested whether the relationship between strong mismatch RTs and posterior alpha MPE effects (mean power in the third Mismatch × Strength cluster in **Fig. 2f**) was mediated by frontal theta effects (mean power in the second Mismatch × Strength cluster in **Fig. 2d**). In all models, frontal theta power, PC3 scores, and posterior alpha power were z-scored within participants and intercept terms were excluded. RTs were log transformed to account for potential outliers and a skewed distribution, then z-scored. An additional exploratory model was tested, examining whether the relationship between mismatch RTs and frontal theta was mediated by posterior alpha.

#### 2.7 Cross-Correlation Analyses

To further characterize the temporal dynamics between the different physiological measures, we conducted cross-correlation analyses on the trial-level 4s-long frontal theta, posterior alpha, and pupil retrieval time series for strong Mismatch trials. Because the pupil time series was downsampled in previous analyses to 100Hz whereas the EEG power time series were downsampled to 250Hz, the pupil time series was upsampled to match the frequency of the EEG power time series. Given the relatively slower dynamics of the pupil, this upsampling should not substantially impact the temporal profile of the pupil time series. A lag window of 1s was defined and the correlation coefficients for different lags were obtained for each pair of trials. Correlation coefficients were averaged across trials for each participant. Significance testing for each lag was performed using one-sampled t-tests and p-values were Bonferroni corrected.

## Results

### 1. Effects of mnemonic prediction errors

Integrating behavior, electrophysiology, and pupillometry, we examined the core hypothesis that mnemonic prediction errors (MPEs) trigger an increase in cognitive control, attention, and arousal (left side of **Fig. 1**). We tested this hypothesis through condition comparisons of prediction errors following strong vs. weak associative encoding. We then explored whether such increases in control, attention, and/or arousal predict subsequent memory for the unexpected/mismatch stimuli.

#### Prediction strength impacts match/mismatch choice behavior

In a combined sample of 101 participants, associative memory *d’* (Z_associative hit_–Z_false alarm_) was higher for strong (3.11±0.93) compared to weak pairings (1.67±0.76) (t_100_=20.988, p<0.001; **Fig. 2c**; see **Supplementary Fig. 2** for data separated by experiment; see **Table 1** for hit and false alarm rates). Consistent with participants retrieving the associate prior to probe onset, probe-locked response times (RT) were relatively rapid, with across-participant mean RT occurring prior to offset of the probe. RTs were faster for strong compared to weak trials (main effect of Strength: β=0.211, CI=[0.202, 0.219], p<0.001), correct than incorrect trials (Accuracy: β=0.200, CI=[0.171, 0.229], p<0.001), and match than mismatch trials (Mismatch: β=0.165, CI=[0.154, 0.175], p<0.001). The RT difference between (a) correct and incorrect trials was larger for strong than weak pairings (Strength × Accuracy: β=−0.167, CI=[−0.194, −0.140], p<0.001); (b) match and mismatch trials was larger for strong compared to weak pairings (Strength × Mismatch: β=−0.126, CI=[−0.141, −0.111], p<0.001); and (c) correct and incorrect trials was larger for match than mismatch trials (Accuracy × Mismatch: β=−0.152, CI=[−0.189, −0.115], p<0.001). The Strength × Accuracy × Mismatch interaction also was significant (β=0.140, CI=[0.096, 0.184], p<0.001). Collectively, these outcomes indicate that: participants followed task instructions, with retrieval of the cued associate tending to begin prior to probe onset, thus establishing conditions for a MPE on mismatch trials; memory was superior following repeated learning (higher *d’* and faster RTs in the strong condition); responses were slower when the probe was a mismatch; and strong MPEs had a greater impact on RTs relative to weak MPEs.

**Table 1:** Hit and false alarm rates.

|  | % Hits (SD) | % False alarms (SD) |
| --- | --- | --- |
| Associative retrieval performance |  |  |
| Strong (4x) | 94.1 (9.9) | 11.1 (13.5) |
| Weak (1x) | 71.0 (18.8) | 17.0 (16.2) |
| Recognition task performance |  |  |
| Experiment 1 |  |  |
| Strong (4x) | 75.6 (16.4) |  |
| Weak (1x) | 76.7 (18.3) |  |
| Familiar item | 43.9 (16.7) |  |
| Novel |  | 15.7 (9.3) |
| Experiment 2 |  |  |
| Strong (4x) | 46.4 (18.7) |  |
| Weak (1x) | 48.6 (20.6) |  |
| Familiar item | 27.6 (15.1) |  |
| Novel |  | 14.2 (8.7) |

#### Strong MPEs signal a need for cognitive control

Frontal theta is posited to mark the engagement of cognitive control and support goal-directed behavior^35^. We examined how MPEs impact frontal theta power, leveraging trial-level regression models to examine the effects of match vs. mismatch probes and condition-level differences between strong vs. weak predictions. We focus here on temporal clusters where there were effects of Strength, Mismatch, and/or an interaction that emerged after probe onset. Because some participants had few to no error trials, models only included correct trials (for models including incorrect trials, see **Supplementary Fig. 3 and Supplementary Tables 1-3**).

Post-probe onset, there was a main effect of Mismatch (0.032 to 0.704s; mean β=0.35, 95% CI=[0.26, 0.44]; p<0.001; **Fig. 2d**), a main effect of Strength (0.504 to 0.872s; β=0.14, CI=[0.07, 0.21]; p<0.001) and a Mismatch × Strength interaction (0.048 to 0.728s; β=−0.35, CI=[−0.46, −0.24]; p<0.001; all p-values for temporal clusters are permutation-corrected, unless indicated otherwise). The interaction reflects a greater difference in frontal theta power between mismatch and match trials on strong compared to weak trials (Mismatch effect size, strong: Cohen’s d=1.28; weak: d=0.85). These data suggest that strong MPEs elicit a greater increase in cognitive control relative to weak MPEs, with these frontal theta power changes primarily preceding responses (**Supplementary Fig. 4a**).

After probe offset (i.e., during the ITI), there was a main effect of Mismatch (0.394 to 0.698s; β=−0.07, CI=[−0.12, −0.02]; p<0.001), a main effect of Strength (0.466 to 0.626s; β=−0.06, CI=[−0.12, 0.00]; p<0.001), and a Mismatch × Strength interaction (0.194 to 0.738s; β=0.11, CI=[0.05, 0.17]; p<0.001); the signs of all three effects were inverted relative to the post-probe onset effects. Indeed, the interaction indicates that the difference in the magnitude of the post-probe offset dip in frontal theta between match and mismatch trials was larger for strong compared to weak trials.

#### Strong MPEs increase attentional engagement

Attentional engagement can be indexed by multiple physiological measures, including a decrease in posterior alpha power^22^. After probe onset, there was a main effect of Mismatch (0.512 to 0.816s; β=−0.13, CI=[−0.18, - 0.08]; p<0.001; **Fig. 2e**), a main effect of Strength (0.504 to 1.480s; β=−0.18, CI=[−0.22, −0.14]; p<0.001), and a Mismatch × Strength interaction (0.536 to 1.376s; β=0.15, CI=[0.10, 0.20]; p<0.001). The interaction indicates that, relative to weak MPEs, strong MPEs evoked greater posterior alpha suppression, putatively marking greater attentional engagement. Moreover, weak matches evoked greater alpha suppression relative to strong matches. Response-locked analyses showed that qualitatively, responses coincided with a drop in posterior alpha, that main effects of Mismatch in posterior alpha were evident prior to responses, and that strength-modulated MPE effects in posterior alpha emerged primarily after responses (**Supplementary Fig. 4b**).

Attentional engagement and arousal also can be indexed by analyses of pupil size. Across 94 participants with suitable pupil data and sufficient trial counts per condition, we found that relative to probe onset, there was a main effect of Mismatch (−0.14 to 2.00s; β=0.16, CI=[0.09, 0.23]; p<0.001; **Fig. 2f**), a main effect of Strength (1.29 to 2.00s; β=0.22, CI=[0.15, 0.29]; p<0.001), and a Mismatch × Strength interaction (0.52 to 2.00s; β=−0.32, CI=[−0.40, −0.24]; p<0.001) (see **Supplementary Fig. 5** for data separated by experiment). Complementing the posterior alpha findings, the interaction indicates that the increase in pupil diameter was larger for strong compared to weak MPEs. This strength-modulated MPE effect specifically emerged after responses (**Supplementary Fig. 4c**).

#### Distinct pupil features reflect mnemonic prediction errors and attentional orienting

We leveraged temporal PCA over pupil data (N=96) to gain further insights into the sequence of cognitive operations that were engaged over the course of the associative retrieval trials and in response to MPEs. As detailed in corresponding Results sections below, the temporal profile of each of six components (**Fig. 3a**) reveal relationships with trial structure, condition, and retrieval behavior, and suggest that the analysis isolated: (a) the effects of pupil-linked arousal at the time of cue presentation on subsequent associative retrieval success (PC2); (b) responses to the strength of the cue (PC2 and PC5); (c) engagement of cognitive effort in support of controlled retrieval (PC5); (d) memory retrieval strength (PC1); (e) responses to probe onset and detection of mismatch stimuli (PC6); (f) responses to MPEs (PC3 and PC4); and (g) post-error orienting responses (PC4). In the following sections, we concentrate on and further delineate specific pupil components that most closely relate to our core hypothesis (for discussion of PC1, PC2, PC5, and PC6, see **Supplementary Fig. 6-8 and Supplementary Tables 4-12**).

**Fig. 3:**
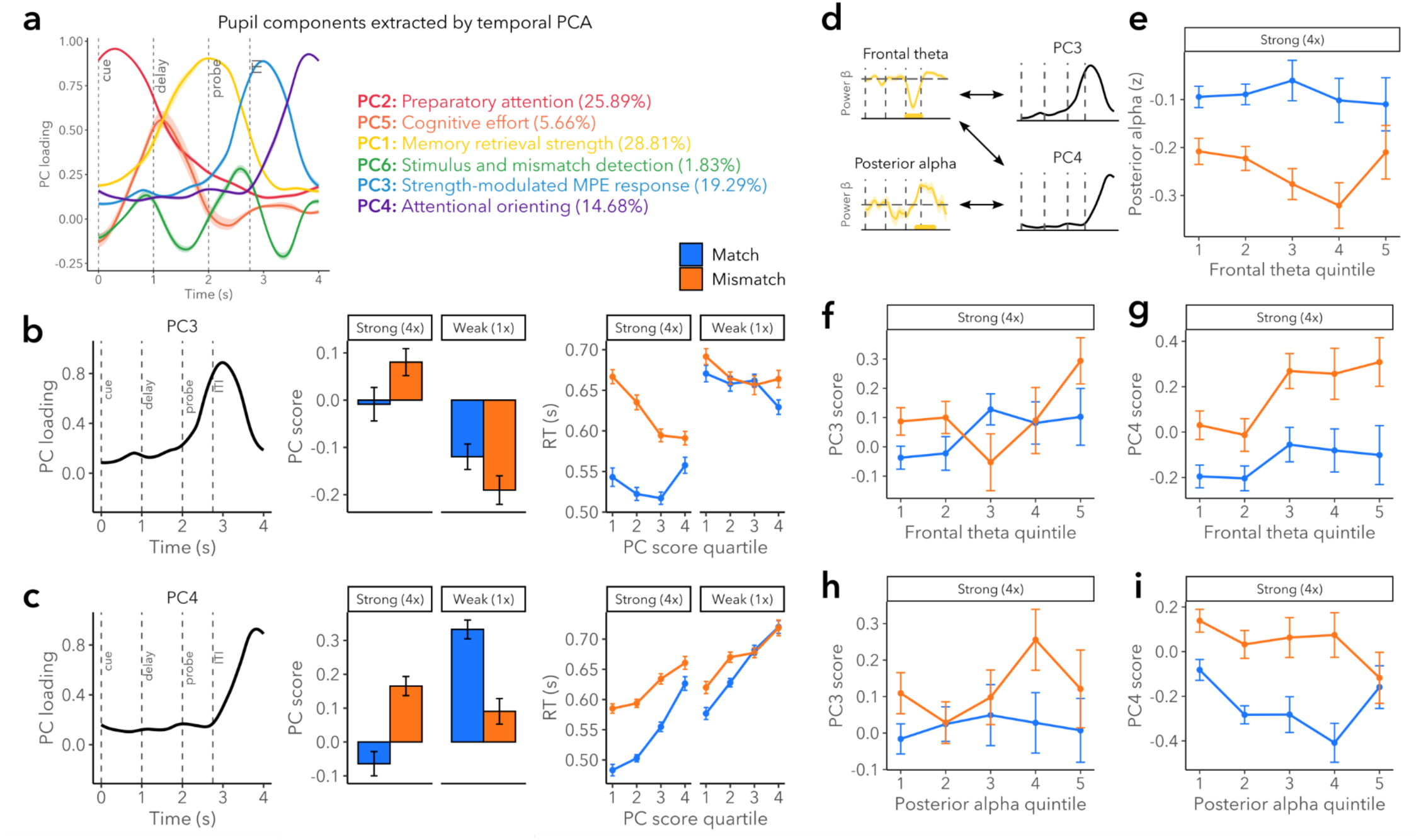
Pupil components sensitive to MPEs and their relationships with frontal theta and posterior alpha. (a) Pupil components extracted by temporal PCA with bootstrapped 95% confidence intervals. PC scores for PC3 (b) and PC4 (c) as a function of match/mismatch conditions and memory strength (middle) and as related to reaction times (right). (d) Schematic of observed relationships among frontal theta, posterior alpha, PC3 scores, and PC4 scores. Time windows over which frontal theta and posterior alpha were averaged in trial-level analyses are underlined in yellow and correspond to those displayed in Fig. 2. (e) Relationship between frontal theta cognitive control signals and attention signals in posterior alpha. (f and g) Relationships between frontal theta cognitive control signals and PC3 (f) and PC4 (g). (h and i) Relationships between posterior alpha MPE-related attention signals and PC3 (h) and PC4 (i). Quartiles and quintiles are shown for visualization purposes only; data were analyzed using continuous measures as described in the Methods.

By relating PC scores to behavior and experimental conditions (memory strength and match/mismatch), we identified two PCs (PC3 and PC4) that were sensitive to MPEs (**Fig. 3b and 3c**). PC3, accounting for 19.29% of variance, peaked 0.99s after probe onset (**Fig. 3b**; 95% CI = [0.97, 1.01]; loading = 0.88, 95% CI=[0.88, 0.90]) and had a full width at half maximum (FWHM) of 1.18s (CI=[1.16, 1.21]). PC4, accounting for 14.68% of variance, peaked towards the end of the trial (**Fig. 3c**; 1.82s after probe onset, CI=[1.81, 1.83]; loading = 0.93, CI=[0.92, 0.93]). To interpret these components and test their sensitivity to MPEs, we examined how the scores for each PC related to prediction strength and match/mismatch conditions; we further assessed how RTs related to PC scores and experimental conditions.

For PC3, there was a Mismatch × Strength interaction (**Table 2**), indicating a strength-sensitive MPE pupil response. Pairwise comparisons showed higher PC3 scores for strong mismatches (correct rejections) than strong matches (hits) (Δ*_Match-Mismatch_*=−0.079, CI=[−0.134, −0.024], t(148.072)=−2.822, p=0.005); for weak pairings, PC3 scores were higher for matches than mismatches (Δ*_Match-Mismatch_*=0.093, CI=[0.034, 0.153], t(197.646)=3.091, p=0.002). Using data from all trials, including misses and false alarms, tPCA extracted highly similar components (r(399)>0.99, p<0.001 for all six PCs); using these component scores, we performed a model comparison to examine whether Accuracy explained a significant amount of variance in PC3 scores. Indeed, an augmented model including effects of and interactions with Accuracy (in addition to Mismatch and Strength) fit PC3 scores better than a model that only included effects of and an interaction between Mismatch and Strength (**χ**^2^(8)=63.90, p<0.001; **Table 3**). The enhanced fit of the augmented model further supports the interpretation that PC3 reflects a strength-sensitive MPE pupil response.

**Table 2:** PC3 scores as a function of Mismatch and Strength.

| Fixed effects | Estimate | CI | p-value | SE | DF |
| --- | --- | --- | --- | --- | --- |
| (Intercept) | 0.002 | [-0.119, 0.123] | 0.979 | 0.061 | 90.970 |
| Mismatch | 0.079 | [0.024, 0.134] | 0.005 | 0.028 | 148.072 |
| Strength | -0.113 | [-0.177, -0.050] | 0.001 | 0.032 | 114.654 |
| Mismatch x Strength | -0.172 | [-0.240, -0.104] | 0.000 | 0.035 | 10236.929 |

**Table 3:** PC3 scores as a function of Mismatch, Strength, and Memory Accuracy.

| Fixed effects | Estimate | CI | p-value | SE | DF |
| --- | --- | --- | --- | --- | --- |
| (Intercept) | 0.034 | [-0.087, 0.155] | 0.577 | 0.061 | 92.514 |
| Mismatch | 0.079 | [0.027, 0.130] | 0.003 | 0.026 | 215.623 |
| Accuracy | 0.021 | [-0.104, 0.146] | 0.743 | 0.064 | 655.824 |
| Strength | -0.107 | [-0.168, -0.046] | 0.001 | 0.031 | 123.938 |
| Mismatch x Accuracy | -0.418 | [-0.601, -0.235] | 0.000 | 0.093 | 3369.869 |
| Mismatch x Strength | -0.176 | [-0.244, -0.107] | 0.000 | 0.035 | 11969.179 |
| Accuracy x Strength | -0.169 | [-0.304, -0.035] | 0.013 | 0.068 | 3106.734 |
| Mismatch x Accuracy x Strength | 0.490 | [0.275, 0.706] | 0.000 | 0.110 | 7003.623 |

Altogether, the RT findings suggest that stronger MPEs (i.e., faster mismatch RTs) evoked larger pupillary responses (right side of **Fig. 2**). This was manifest in there being a Mismatch × Strength × PC3 score interaction (**Table 4**), with post-hoc analyses showing that higher PC3 scores for strong mismatches were coupled with faster responses (estimated slope=−0.046, CI=[−0.061, −0.031], t(306.238)=−6.219, p<0.001), whereas RTs and PC3 scores were unrelated for strong matches (estimated slope=0.009, CI=[−0.003, 0.021], t(152.586)=1.463, p=0.145); the difference between these slopes was significant (match – mismatch slope=0.055, CI=[0.038, 0.072], t(4019.950)=8.196, p<0.001). Furthermore, RTs and PC3 scores were more strongly related for strong compared to weak mismatches (strong – weak slope=−0.032, CI=[−0.053, −0.012], t(5920.054)=−4.134, p<0.001). Together, these results further support the interpretation that PC3 is a strength-sensitive MPE component that is sensitive to differences in prediction strength at both the condition-level (strong vs. weak) and trial-level (e.g., trial-level strong mismatch RTs).

**Table 4:**
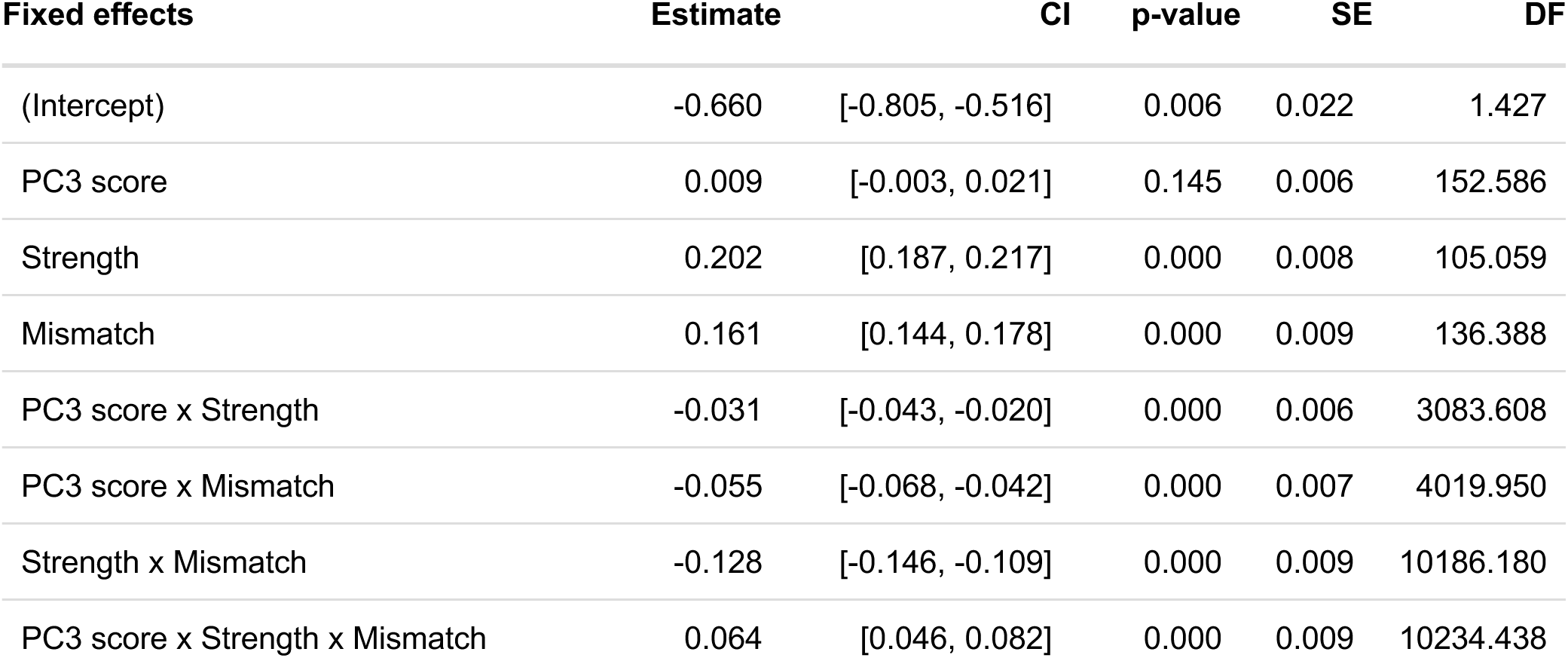
PC3 scores and RTs.

A different pattern was observed for weak trials, with only a trend-level relationship between PC3 scores and RTs for mismatches (estimated slope=−0.014, CI=[−0.029, 0.001], t(341.456)=−1.777, p=0.077), a negative relationship for matches (estimated slope=−0.022, CI=[−0.035, −0.010], t(188.098)=−3.450, p=0.001), and no significant difference between these slopes (match – mismatch slope=−0.009, CI=[−0.027, 0.010], t(4473.238)=−1.243, p=0.599). These outcomes are consistent with the behavioral findings that MPEs were experienced primarily on strong but not weak trials (**Fig. 2**). In addition, these findings indicate that within the strong condition, when stronger mnemonic predictions (as indexed by shorter strong mismatch RTs) were violated, larger pupillary responses were evoked.

Similar to PC3, PC4 responses displayed strength-sensitive MPE modulation (**Fig 3c**; **Table 5**). A Mismatch × Strength interaction revealed a larger difference between mismatch and match scores in the strong than the weak condition; strong mismatch scores were higher than strong match scores (Δ*_Match-Mismatch_*=−0.275, CI=[−0.327, −0.224], t(10260.360)=−10.486, p<0.001) and weak mismatch scores were lower than weak match scores (Δ*_Match-Mismatch_*=0.202, CI=[0.145, 0.258], t(10269.580)=6.947, p<0.001). PC4 scores from the dataset including error trials were better fit by a model that additionally included effects of and interactions with Accuracy (**χ**^2^(8)=213.83, p<0.001); moreover, component scores were highest for strong error trials (**Table 6**; Accuracy effect; Accuracy × Strength).

**Table 5:**
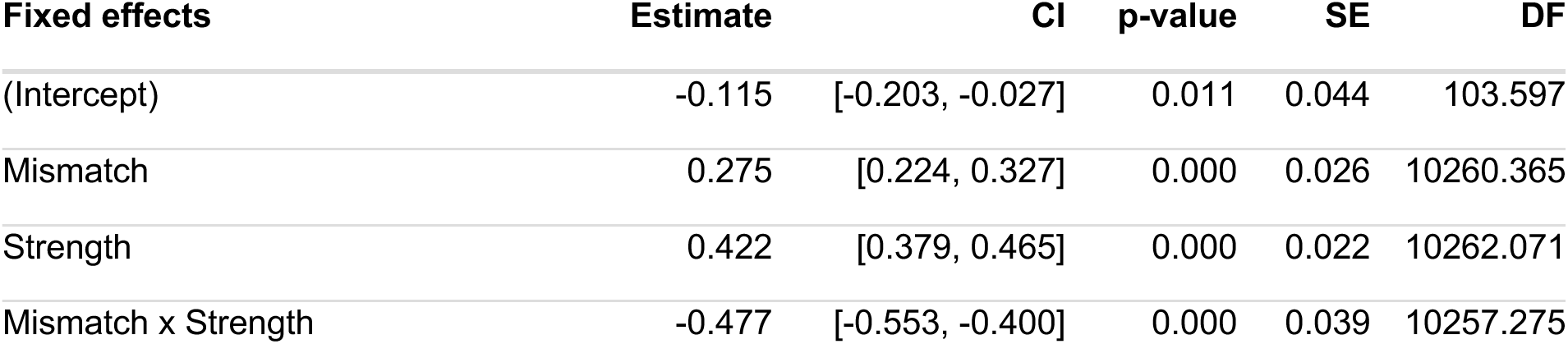
PC4 scores as a function of Mismatch and Strength.

**Table 6:**
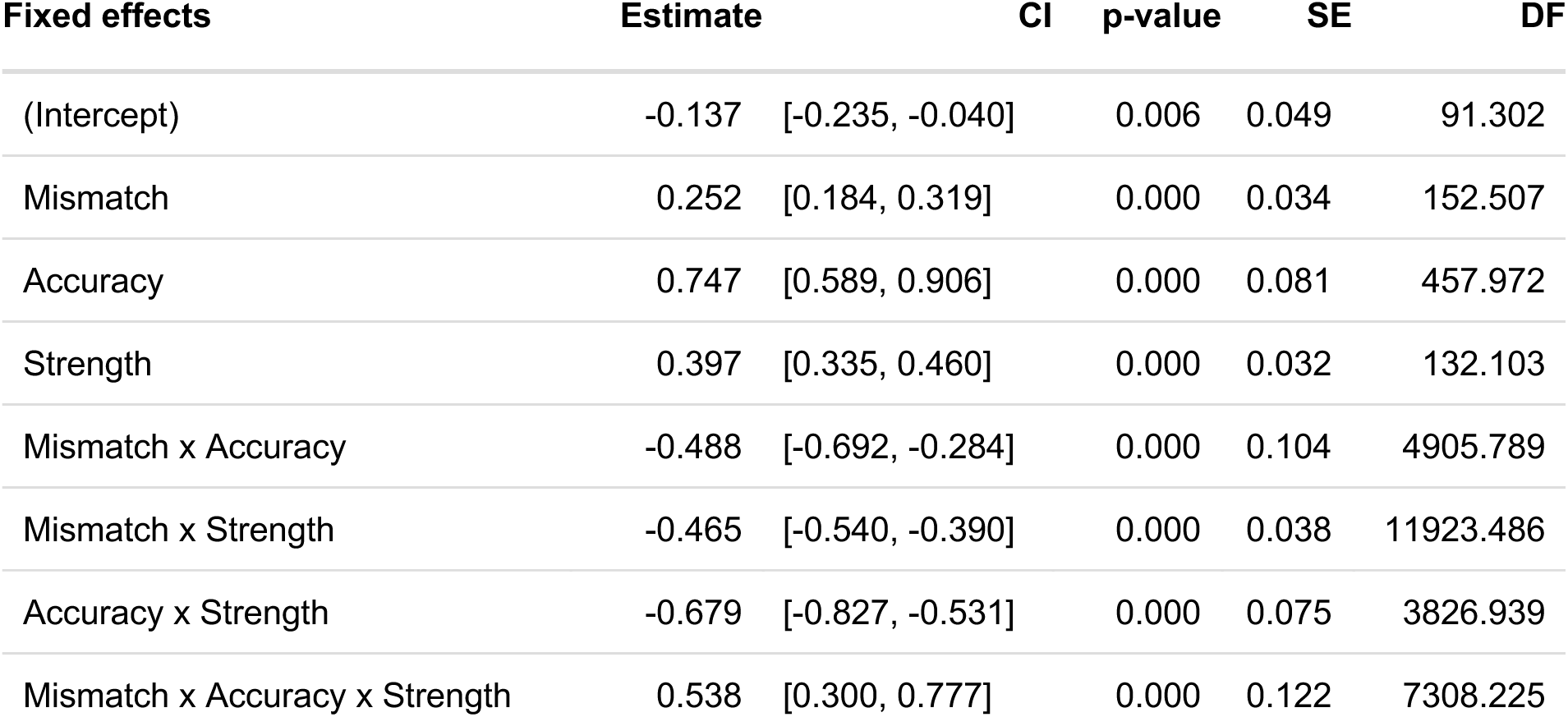
PC4 scores as a function of Mismatch, Strength, and Memory Accuracy.

The RT model showed that weaker mnemonic predictions (or greater difficulty in correctly retrieving memories), as indexed by longer RTs, were associated with higher PC4 scores (**Table 7**; effect of PC4 score). The relationship between PC4 scores and RTs was stronger for weak compared to strong trials (PC4 score × Strength) and for match compared to mismatch trials (PC4 score × Mismatch). There was no three-way interaction (PC4 score × Mismatch × Strength). As such, more effortful memory retrieval, manifesting as slower RTs and/or erroneous responses, elicited larger delayed pupillary responses.

**Table 7:**
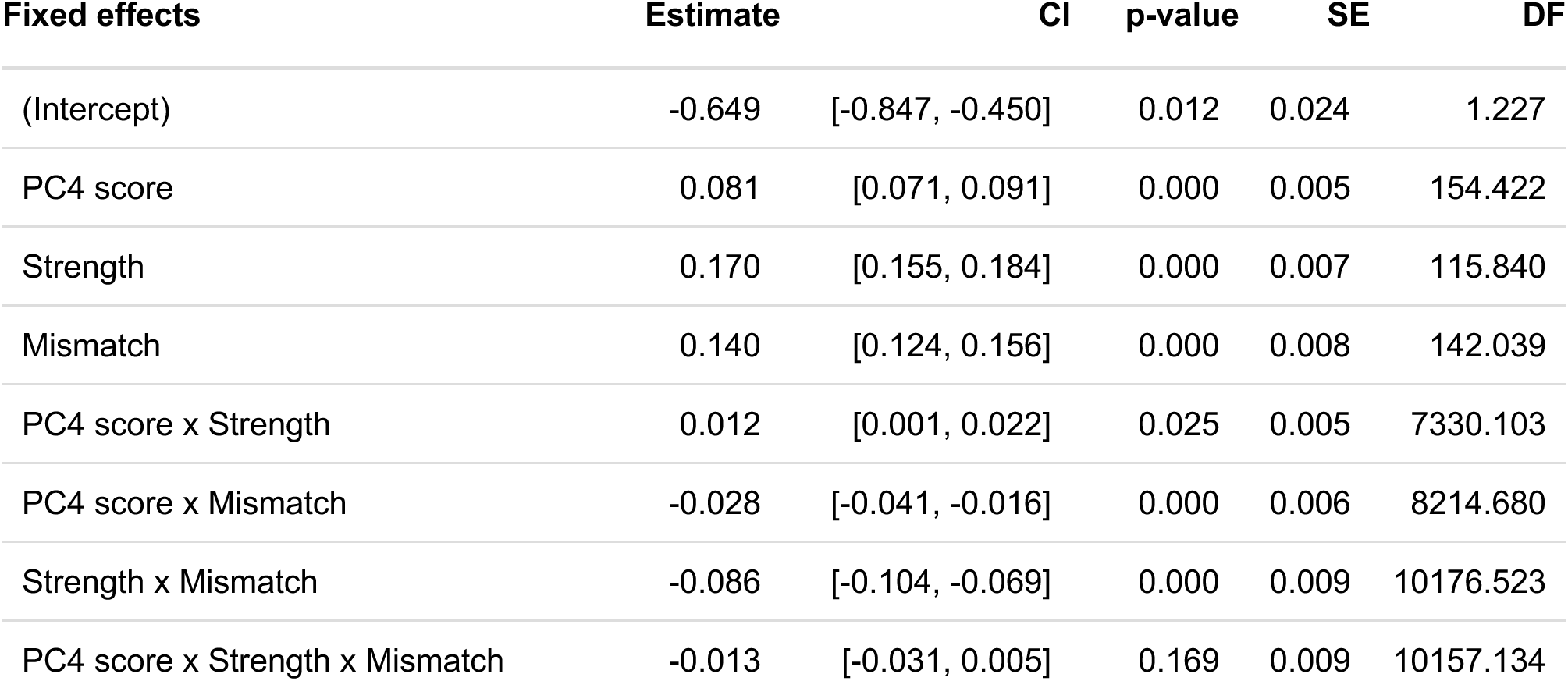
PC4 scores and RTs.

Together, these results suggest that PC4, which peaked after PC3 and after responses, reflects post-MPE cognitive processing and perhaps a change in attentional engagement. PC4 appears to primarily reflect an orienting response to errors – not only strong erroneous predictions, but also erroneous memory decisions. This component is sensitive to the amount of effort (assayed by RTs) exerted to generate those predictions and the strength of those errors (strong vs. weak). Elevated PC4 scores on weak hits, where no error was made, may instead reflect retrieval practice and thus internally oriented attention. On these trials, when cue-elicited retrieval may have been weaker, the probe may have provided additional support for pattern completion of the learning episode for that association, and this engagement in memory retrieval may have elicited pupil dilation (c.f., Strength effect in **Fig. 2f**). Overall, PC4 may therefore reflect an attentional orienting response that, depending on the relative success of memory retrieval and probe identity, may direct attention internally or externally.

PC3 and PC4 scores were differentially associated with RTs, with PC3 showing an inverse association during strong MPEs and PC4 showing positive associations in all conditions. This differential relationship suggests for stronger MPEs (i.e., strong MPEs that were responded to quickly), higher PC3 scores may reflect more rapid resolution of conflict and lower PC4 scores may therefore reflect less attentional orienting. In other cases, when PC4 scores are high, conflict resolution may have taken longer and subsequent orienting responses may not resolve before the onset of the subsequent trial. As such, PC3 and PC4 may have differential effects on cognition and behavioral performance on subsequent trials.

#### Summary

Following probe onset, detection of a mismatch may elicit engagement of cognitive control (i.e., an increase in frontal theta) to facilitate comparison of mnemonic and sensory evidence. Sustained attention to mnemonic evidence and evidence accumulation during the comparison process may drive an anticipatory pupil response up until the decision; the amplitude of this response may ultimately be impacted by the magnitude of the MPE (PC3). Upon completion of the trial, after a response is made, additional cognitive processes may lead to subsequent changes in pupil size (PC4) that reflect additional changes in attention/arousal.

We next investigated whether, and if so, how control processes interacted with attention/arousal by examining relationships between MPE-related frontal theta, posterior alpha, and pupil responses.

### 2. Relationships between control, attention, and arousal

#### Frontal theta predicts pupillary responses, but not posterior alpha

Altogether, our behavioral, electrophysiological, and pupillary findings provide converging evidence that stronger MPEs were experienced in the strong condition relative to the weak condition. Accordingly, we examined the relationship between MPE-related impacts on frontal theta, posterior alpha, and pupil, focusing on strong trials and the effects of matches and mismatches.

Changes in attention and arousal following MPEs may be elicited by error detection alone or by a separate signal that more control is needed. To investigate this question, we examined whether putative cognitive control signals, assayed by increased frontal theta power, predicted changes in posterior alpha and pupil size (**Fig. 3d**). First, we observed no relationship between MPE-evoked frontal theta (second Mismatch × Strength cluster) and posterior alpha (third Mismatch × Strength cluster) (**Fig. 3e**; main effect of frontal theta: β=0.010, CI=[−0.012, 0.032], p=0.369; frontal theta × Mismatch interaction: β=−0.023, CI=[−0.056, 0.011], p=0.182]). Second, while higher frontal theta on strong mismatches was associated with higher decision-related pupil components — both immediate (**Fig. 3f**; PC3: main effect of frontal theta: β=0.050, CI=[0.008, 0.092], p=0.020) and lagged (**Fig. 3g**; PC4: main effect of frontal theta: β=0.054, CI=[0.005, 0.104], p=0.033) — the strength of this coupling did not differ between matches and mismatches for PC3 (frontal theta × Mismatch interaction for PC3: β=0.001, CI=[−0.056, 0.059], p=0.966) or PC4 (frontal theta × Mismatch interaction for PC4: β=0.024, CI=[−0.055, 0.103], p=0.544]). Finally, frontal theta predicted mean pupil diameter (Mismatch × Strength cluster) (**Supplementary Fig. 9a**; main effect of frontal theta: β=0.046, CI=[0.007, 0.084], p=0.022), with no difference in the strength of this relationship between mismatch and match trials (frontal theta × Mismatch interaction (β=0.038, CI=[−0.021, 0.097], p=0.212).

Together, these results indicate that the need for cognitive control, as assayed by frontal theta, was associated with immediate and lagged pupil-linked increases in attention and arousal (i.e., PC3, PC4, and mean pupil diameter). Although frontal theta was higher for strong mismatches than strong matches (**Fig. 3d**), the strength of the relationship between frontal theta and pupillary responses did not differ as a function of MPE detection. Increases in attention as assayed by posterior alpha were independent of frontal theta control signals. Thus, these outcomes suggest that some changes in attention following MPEs are elicited by error detection alone (i.e., posterior alpha), whereas other changes in attention/arousal (i.e., pupil) are associated with control signals.

#### Posterior alpha couples with lagged pupillary orienting responses, but not immediate decision-related pupil responses

We next examined whether MPE-driven changes in posterior alpha were associated with pupil-linked attention and arousal responses. There was no relationship between posterior alpha power and PC3 scores (**Fig. 3h**; main effect of posterior alpha: β=−0.005, CI=[−0.077, 0.067], p=0.894; PC3 score × Mismatch: β=0.017, CI=[−0.108, 0.141], p=0.793). By contrast, PC4 scores and mean pupil diameter were negatively coupled with posterior alpha power (**Fig. 3i**; main effect of posterior alpha on PC4 scores: β=−0.155, CI=[−0.229, −0.080], p<0.001; **Supplementary Fig. 9b**; main effect of posterior alpha on mean pupil: β=−0.107, CI=[−0.182, −0.032], p=0.007), indicating that greater posterior alpha suppression (a marker of attentional engagement) predicted larger pupillary orienting responses (a marker of increased arousal and/or attention). There was no PC4 score × Mismatch interaction (β=−0.053, CI=[−0.192, 0.086], p=0.452) nor was there a mean pupil × Mismatch interaction (β=0.028, CI=[−0.112, 0.168], p=0.687). As such, greater attentional engagement following memory decisions, as assayed by decreases in posterior alpha, was followed by stronger pupillary orienting responses.

Together, these findings suggest that the magnitude of alpha suppression following memory decisions is associated with lagged orienting pupillary responses, but not immediate decision-related pupillary responses. In other words, the initial MPE-evoked changes in posterior alpha and pupil size (PC3) are independent of each other.

#### Strong MPE-evoked increases in cognitive control do not explain increases in attention and arousal

While the preceding outcomes suggest that attention, arousal, and control processes are triggered at least partially independently by MPEs, we conducted multiple mediation analyses to further test whether the dynamics of observed cognitive and neural changes following strong MPEs are consistent with or challenge our hypothesis that changes in cognitive control partially drive increases in attention and arousal. In these analyses, we focused on the dynamics in response to strong MPEs (strong mismatch trials) and used RTs as a proxy measure for the trial-level magnitude of the MPE (with shorter RTs reflecting stronger predictions and thus stronger MPEs). Pupil and posterior alpha effects were modeled separately; in each model, we tested whether the trial-level relationships between MPE magnitude (RTs) and changes in attention and arousal (pupil PC3/posterior alpha) were mediated by increases in cognitive control (frontal theta) (**Fig. 4a**). In the pupil model (**Fig. 4b**), the credible interval for the effect of RTs on frontal theta included zero, providing no strong evidence for this relationship (*a*=−0.052, 95% credible interval (CrI)=[−0.137, 0.034]). There was strong evidence for an effect of frontal theta on PC3 scores (*b*=0.070, CrI=[0.002, 0.138]) and insufficient evidence for frontal theta mediating the relationship between RTs and immediate pupil responses (*a*b*=0.005, CrI=[−0.010, 0.027]). There was strong evidence for a direct effect of RTs on PC3 scores (*c’*=−0.214, CrI=[−0.295, −0.133]). In the posterior alpha model (**Fig. 4c**), there was no strong evidence for an effect of RTs on frontal theta (*a*=−0.052, CrI=[−0.138, 0.033]), as in the pupil model. There was strong evidence that higher frontal theta predicted lower posterior alpha (*b*=−0.082, CrI=[−0.146, - 0.018]) and insufficient evidence for frontal theta mediating the relationship between RTs and posterior alpha (*a*b*=0.009, CrI=[−0.004, 0.029]). There was strong evidence for RTs negatively predicting posterior alpha (*c’*=−0.124, CrI=[−0.189, −0.058]). Together, these outcomes suggest that increases in attention and arousal following strong MPEs are not well-explained by increases in cognitive control. Yet, given that these exploratory analyses may be underpowered, caution is warranted in interpreting these null findings.

**Figure 4.**
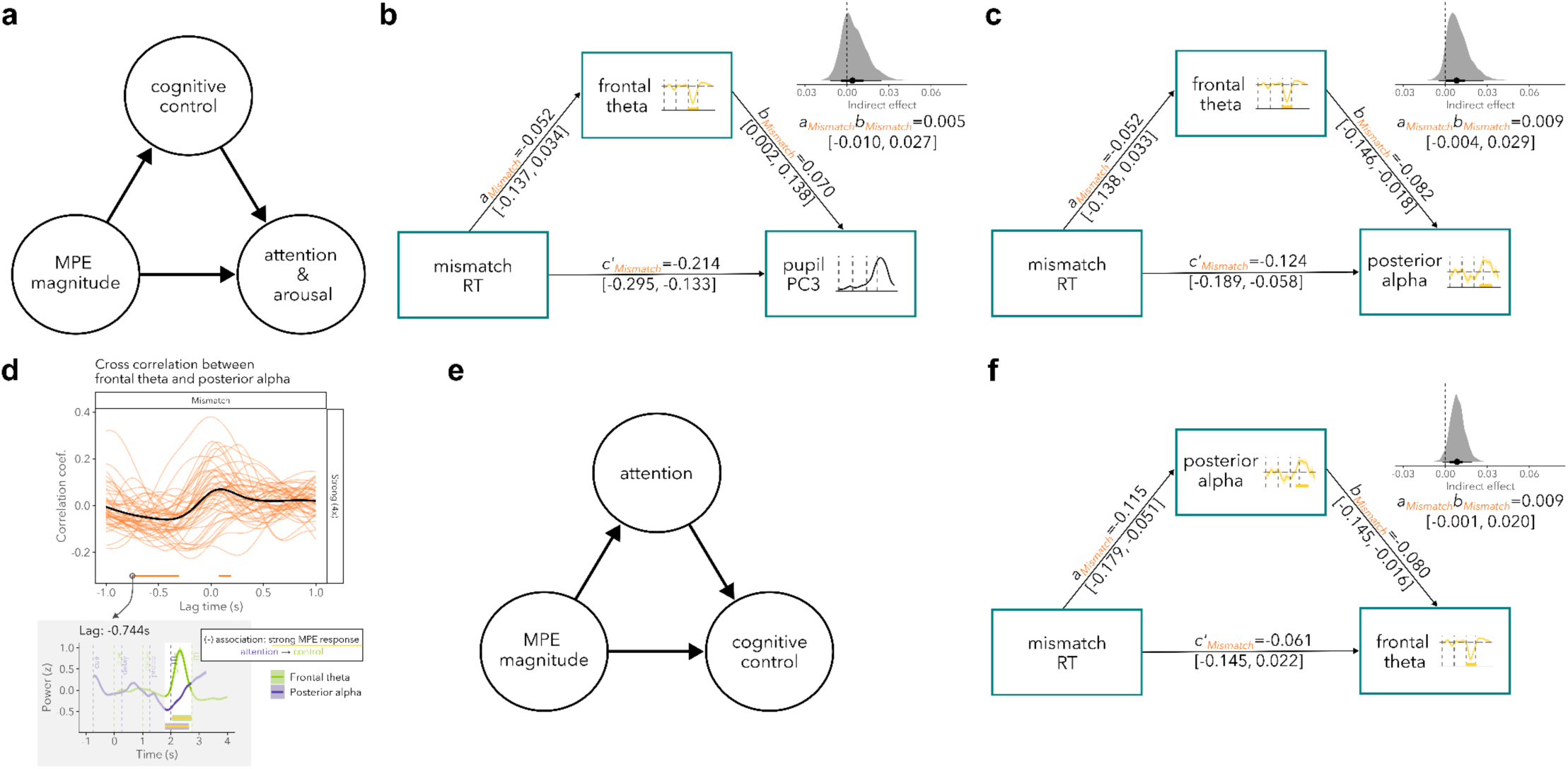
Dynamics between cognitive control, attention, and arousal in response to strong MPEs. (a) Schematic of hypothesized relationships: stronger MPEs increase attention/arousal and this effect was hypothesized to be partially explained by an increase in cognitive control. (b) A multivariate mediation analysis testing the model diagrammed in (a) with pupil PC3 as the measure of attention/arousal. (c) A multivariate mediation analysis testing the model diagrammed in (a) with posterior alpha as the measure of attention. (d) Cross-correlation analysis showing the correlation between frontal theta time series and lagged posterior alpha time series on strong MPE trials. (e) An alternative model: changes in cognitive control following MPEs may be explained by changes in attention, as assayed by posterior alpha. (f) A multivariate mediation analysis testing the model in (e), examining whether posterior alpha mediates the relationship between RTs and frontal theta. Paths in mediation models are labeled with the mean of the posterior and 95% credible intervals. Gray histograms accompanying mediation model diagrams depict the posterior distribution for the indirect effect; black points indicate posterior medians and horizontal lines indicate the 66% and 95% highest density intervals.

#### Posterior alpha responses to strong MPEs precede increases in frontal theta

To develop further insights into the temporal relationships among the neurocognitive measures, we explored their temporal relationships using cross-correlational analyses with a lag window of 1s. For frontal theta and posterior alpha, there were two significant clusters with dissociable lag patterns. Interpretation of these clusters was guided by the direction of the relationship between frontal theta and posterior alpha and how this direction compared with prior analyses; after identifying candidate cognitive processes associated with a given lag cluster, we then used the lag time to make inferences about which cognitive process was leading or lagging.

There was a cluster with a negative mean correlation and with posterior alpha leading (−0.744 to −0.312s, mean r=−0.053, CI=[−0.055, 0.052]). The negative relationship between frontal theta and posterior alpha – greater control and more attention – may primarily reflect cognition in response to the strong MPE, given the differential sign of the primary Mismatch × Strength interactions in **Fig. 2d** and **Fig 2e**^a^. The negative lag suggests that an increase in attention, as assayed by posterior alpha, may precede an increase in cognitive control as indexed by frontal theta. This interpretation is consistent with the outcomes in the response-locked trial-level regression models (**Supplementary Fig. 4a** and **Supplementary Fig. 4b**), which showed that qualitatively, main effects of Mismatch emerged for posterior alpha before main effects of Mismatch emerged for frontal theta. While suggestive, future work should more directly examine the possibility that posterior alpha-related changes in attention precede and perhaps drive changes in cognitive control as assayed by frontal theta. The second significant cluster may be associated with retrieval-related processes and is discussed in the Supplement (**Supplementary Fig. 10**). Cross-correlation analyses of pupil with frontal theta and pupil with posterior alpha did not reveal significant correlation clusters.

#### Exploratory analyses examining whether changes in posterior alpha explain increases in frontal theta

Given the finding that posterior alpha responses may precede frontal theta responses to strong MPEs, we tested an alternative mediation model in an exploratory analysis to assess the possibility that changes in cognitive control are partially explained by changes in attention as indexed by posterior alpha (**Fig. 4e**). There was strong evidence for an effect of RTs on posterior alpha (**Fig. 4f**; *a*=−0.115, CrI=[−0.179, −0.051]) and strong evidence for an effect of posterior alpha on frontal theta (*b*=−0.080, CrI=[−0.145, −0.016]). However, there was insufficient evidence for an indirect effect (*a*b*=0.009, CrI=[−0.001, 0.020]). There was also insufficient evidence for a direct effect of RTs on frontal theta (*c’*=−0.061, CrI=[−0.145, 0.022]), as in the earlier mediation models (**Fig. 4b-c**). Bayesian leave-one-out cross-validation showed that the difference in model fit for alpha as a mediator (**Fig. 4c**; ELPD=−2842.07) and theta as a mediator (**Fig. 4f**) was minimal (ΔELPD=−0.82±0.41). As such, the current findings do not provide conclusive evidence for or against the idea that control signals modulate attention, or vice versa.

#### Summary

In sum, strong MPEs evoked changes in frontal theta (**Fig. 2d**), posterior alpha (**Fig. 2e**), and pupil diameter (**Fig. 2f**). While the changes in frontal theta and posterior alpha were not significantly associated with each other following strong matches and showed no interaction with Mismatch, there was suggestive evidence from the mediation models that they are negatively coupled in response to strong MPEs – with greater control predicting more attention. Changes in frontal theta and posterior alpha were both associated with the magnitude of late pupil responses (PC4). Initial decision-related pupil responses (PC3) were associated with frontal theta but not posterior alpha. These relationships did not differ between Match and Mismatch, suggesting that coupled changes in attention/arousal and control during/following memory decisions are not specific to MPEs. Differences in the patterns of relationships for pupil components and mean pupil diameter highlight the increased sensitivity offered by temporal PCA in identifying pupil responses to varying cognitive processes engaged at different timepoints. Critically, these outcomes highlight the distinct temporal profiles of the relationship between control and attention/arousal: pupillary assays of increased attention/arousal evoked by memory decisions are associated with cognitive control signals (i.e., PC3, PC4, mean pupil), whereas increases in attention as assayed by posterior alpha only relate to later pupil responses (i.e., PC4 but not PC3 or mean pupil). Moreover, cross-correlation analyses suggested that upregulation of attention, as assayed by posterior alpha, precedes changes in cognitive control. Although this outcome suggests that posterior alpha changes may partially explain changes in frontal theta, mediation analyses did not provide strong evidence for control signals upregulating attention or attention increases upregulating control signals.

### 3. Learning from MPEs

#### MPEs similarly enhance recognition of strong and weak mismatches

To investigate whether prediction strength modulates MPE-driven learning, we examined recognition memory for mismatch probes on a subsequent surprise memory test administered after a ∼5-min delay (Experiment 1; E1) or a 48-hr delay (Experiment 2; E2). A 48-hr delay was introduced in Experiment 2 to address the possibility of ceiling recognition performance with the ∼5-min delay in Experiment 1 and to test whether the effects of prediction strength on learning specifically emerge following overnight consolidation. Contrary to our predictions, subsequent recognition (*d’;* **Fig. 5a**) was comparable for strong (E1: 1.81±0.59; E2: 1.02±0.52) and weak mismatch probes (E1: 1.86±0.65; E2: 1.08±0.60) (E1: β=−0.05, CI=[−0.31, 0.21], p=0.89; E2: β=−0.06, CI=[−0.34, 0.21], p=0.852). Recognition of strong (E1: β=0.94, CI=[0.68, 1.20], p<0.001; E2: β=0.52, CI=[0.25, 0.80], p<0.001) and weak (E1: β=0.99, CI=[0.73, 1.25], p<0.001; E2: β=0.58, CI=[0.31, 0.86], p<0.001) mismatch probes was higher than that of familiar items (E1: 0.87±0.48; E2: 0.50±0.45), suggesting that MPEs enhanced memory but stronger MPEs do not further boost recognition of prediction-violating items. See **Table 1** for hit and false alarm rates and **Supplementary Fig. 11** for recognition RTs.

**Fig. 5:**
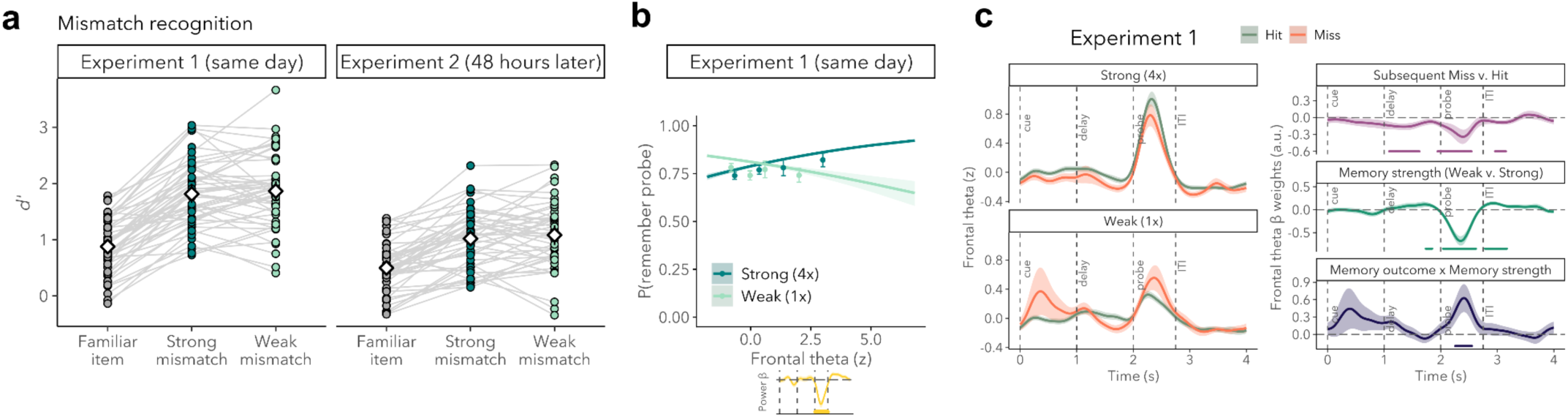
MPE-driven increases in frontal theta by strong MPEs enhance learning. (a) Behavioral performance in the surprise recognition task in Experiment 1 and Experiment 2. (b) Higher frontal theta during strong MPEs is associated with a higher likelihood of subsequently remembering the probe ∼5 min later (Experiment 1). Points indicate binned values for visualization; smoothed lines reflect model estimates. (c) Trial-level regression models for frontal theta modeling the effects of Subsequent Memory, Strength, and Subsequent Memory × Strength interactions in Experiment 1.

#### Frontal theta during strong MPEs enhances learning

To understand whether MPE-driven changes in cognitive control, attention, and/or arousal support learning, we examined whether electrophysiological and pupillary responses to MPEs predict subsequent memory for mismatch probes. In Experiment 1, there was a main effect of mean MPE-related frontal theta (the second Mismatch × Strength cluster) on subsequent memory (odds ratio (O.R.) = 1.186 [1.006, 1.398], p=0.042; **Fig. 5b**; right side of **Fig. 1**) and a significant frontal theta × Strength interaction (O.R.=0.745, [0.597, 0.930], p=0.009), indicating that this relationship was attenuated for weak mismatch probes. While frontal theta during strong MPEs partially accounted for recognition memory of strong mismatch probes, frontal theta during weak MPEs did not explain memory for weak mismatch probes. By contrast, MPE-associated changes in posterior alpha and pupil were not associated with subsequent memory (see **Supplementary Fig. 12**).

Trial-level regression models of frontal theta (**Fig. 5c**) further revealed main effects of Subsequent Memory from 1.088 to 1.624s (β=−0.15, CI=[−0.25, −0.05]; p<0.001), 1.944 to 2.544s (β=−0.24, CI=[−0.42, −0.06]; p<0.001), and 2.960 to 3.152s (β=−0.09, CI=[−0.15, −0.02]; p<0.001) after cue onset, such that frontal theta was higher for subsequent hits than subsequent misses. There was a significant Subsequent Memory × Strength interaction from 0.256 to 0.552s after probe onset (β=0.54, CI=[0.11, 0.98]; p<0.001) whereby the difference in power between hits and misses was larger for strong compared to weak probes. This analysis revealed that frontal theta partially accounts for subsequent memory regardless of strength condition during select time windows during the delay period and following the probe period. In other words, subsequent memory for strong and weak mismatch probes is partially predicted by the level of cognitive control engaged during mnemonic prediction generation and by increases in cognitive control following a mismatch. Greater frontal theta immediately elicited by strong MPEs during the probe period did not additionally enhance memory for strong mismatch probes above that of weak mismatch probes. Collectively, these findings suggest that cue-evoked and delay-period frontal theta, strength-modulated increases in frontal theta following MPEs, and the magnitude of the post-peak frontal theta responses enhanced learning of probes that violated mnemonic predictions.

## Discussion

Features of present experiences often overlap with those of the past. Shared features allow for aspects of current experience to cue memories and enable knowledge about how similar events unfolded in the past to inform predictions about ongoing experience. When ongoing experience unexpectedly diverges from mnemonic predictions, multiple neurocognitive processes may be elicited, fostering adaptation to novel circumstances and facilitating more accurate predictions in the future. Drawing on behavioral, electrophysiological, and pupillary assays, the current study revealed that strong MPEs elicit increases in cognitive control, attention, and arousal, with potential impacts on later cognition and new learning (for a summary, see **Fig. 1**).

### Neurocognitive impacts of MPEs are strength-sensitive

Across three physiological measures – frontal theta, posterior alpha, and pupil size – MPE-related responses were larger when predictions were strong compared to weak. These findings are consistent with the idea that responses to prediction errors are modulated by how *unexpected* the error is^16,17,61^ and relatedly, how much uncertainty there is when making a decision^62^. In the current study, weak predictions may give rise to less certainty about an upcoming event (i.e., the probe) such that weak MPEs may be considered an outcome of an *expected* degree of uncertainty following a weak memory cue. On the other hand, more strongly cued predictions – evidenced in the current study by better memory and faster RTs for strong vs. weak pairs – may elicit lower levels of uncertainty; a MPE in this case would be more unexpected and warrant increasing attention to facilitate model updating. Theories of expected and unexpected uncertainty implicate different neuromodulators in tracking different types of uncertainty, with LC norepinephrine release posited to track unexpected uncertainty^17^. The present pupil size findings, when interpreted as an indirect assay of noradrenergic LC activity^14,15^, are consistent with this theory. Mnemonic prediction strength critically modulates demands on control, attention, and arousal.

### Attention and control dynamics following strong MPEs

Increases in attention and arousal following strong MPEs, as assayed by posterior alpha and pupil, were not well explained by changes in cognitive control. Exploratory cross-correlation analyses showed that increases in attention in response to strong MPEs, as assayed by posterior alpha, may precede increases in cognitive control. While this result suggests that changes in attention may be elicited first and subsequently upregulate cognitive control, an exploratory formal test of this hypothesis did not provide strong evidence for posterior alpha mediating the effects on frontal theta. Given that these analyses were restricted to strong MPE trials, they may not have been sufficiently powered to detect indirect effects or to detect meaningful differences between these models. Of note, while longer mean RTs for mismatches compared to matches are a behavioral marker of a MPE, within-condition differences in RTs (i.e., between mismatch trials) may reflect more subtle differences in MPE magnitude, with shorter RTs reflecting stronger predictions and thus stronger MPEs. Strong mismatch RTs were used in the mediation models as a proxy measure of MPE magnitude; this estimate may be noisy because RTs in this experiment are likely sensitive to factors independent of mnemonic prediction strength (e.g., preparatory attention (**Supplementary Fig. 7**) or memory strength of the mismatch probe). This limitation may have additionally reduced sensitivity to detecting indirect effects. Whether upregulation of attention indeed emerges prior to increases in control, and whether upregulation of attention influences how much control is engaged, remain open questions to be more directly tested in future research.

### Memory modes following mnemonic predictions: Open questions and future directions

Notably, for pupil size, the direction of the effect following weak MPEs was the inverse of that observed following strong MPEs, such that there was greater pupil dilation for weak matches than weak mismatches. We propose that this difference may reflect attention being oriented internally to retrieved contents after a weak mnemonic prediction is confirmed, and that increased pupil dilation following weak matches may reflect the probe facilitating pattern completion of the learning episode for the weak association. These findings suggest that the engagement of retrieval^63,64^ or encoding modes^1,65^ following confirmation or violation of mnemonic predictions may be dependent on prediction strength. However, whether increased pupil dilation following the confirmation of weak mnemonic predictions indeed reflects retrieval requires investigation of the mental contents and targets of increased attention following weak matches.

### Pupil responses to MPEs: Temporal dynamics and neural interactions

Pupil responses during associative retrieval trials indexed multiple cognitive processes. As observed by others^7,8^, temporal PCA revealed multiple pupil responses sensitive to task structure. Moreover, we observed strong coupling between PC scores and trial-level behavioral responses, which enabled interpretation of individual components as reflections of distinct cognitive processes. With respect to MPEs, temporal PCA yielded finer-grained insights into the temporal dynamics of pupillary responses. In particular, this analysis isolated two temporally distinct pupil responses to MPEs: a more immediate strength-modulated prediction error response (PC3) and a later attentional orienting response (PC4). By obtaining trial-level PC scores, as opposed to condition-and subject-level scores as in prior work^7,8^, we were able to relate the multifaceted pupil responses to electrophysiological signals on a trial-by-trial basis. These trial-level analyses offered greater sensitivity to relationships that the pupil has with frontal theta markers of cognitive control and posterior alpha markers of attention. These analyses highlighted that PC3 scores were predicted by engagement of cognitive control (frontal theta) during memory decisions regardless of prediction error. PC4, like PC3, was associated with cognitive control (frontal theta) during memory decisions. In addition, PC4 scores varied with the degree of posterior alpha suppression evoked following memory decisions. Altogether, these findings highlight that initial MPE-related changes in attention and/or arousal assayed separately by pupil and posterior alpha are distinct, but that over time, these measures converge and may subsequently reflect a unitary focus of attention.

### Beyond mismatch signals in PFC: Potential interactions with the hippocampus, VTA/SN, and LC

The hippocampus is thought to compute MPEs^66–69^, with the CA1 subfield being anatomically well-suited for mismatch detection given that it is a convergence zone for mnemonic predictions from hippocampal subregion CA3 and sensory input from entorhinal cortex^70–73^. Functional neuroimaging studies provide support for CA1 as a mismatch “detector” in humans^21,47,74^. Moreover, following MPE signals from the hippocampus, LC, or other regions, learning rates in the hippocampus are posited to increase^70^. Such increases may be underpinned, at least in part, by a biasing of hippocampal functional connectivity towards an encoding-favored configuration^65^, which may enhance the integration of bottom-up sensory information during hippocampal encoding computations^75^. Separately, increases in hippocampal theta power^30,76^ may increase the likelihood of encoding novel information into memory^46^. Finally, dynamic causal modeling of magnetoencephalography (MEG) data suggests that mismatch detection in the hippocampus, as assayed by theta power, may be driven by theta-related signals from ventromedial prefrontal cortex (vmPFC)^30^. Hippocampal theta, which may be challenging to detect with scalp EEG^77^, could modulate or be modulated by cognitive control and attention signals that are often assayed using scalp EEG.

Drawing on frontal theta, the current study characterized the temporal profile of cognitive control following strong MPEs. Additional and more direct corroboration of prior MEG findings that vmPFC theta drives hippocampal theta^30^ is needed to resolve questions about MPE-driven hippocampal-PFC interactions and their relative timing. Are MPEs detected first by PFC or by hippocampus (e.g., within subfield CA1)? Does prediction strength modulate the magnitude of frontal and hippocampal theta responses similarly? Alternatively, does the strength of MPEs modulate hippocampal theta linearly and frontal theta nonlinearly, such that frontal theta primarily responds to strong MPEs?

Hippocampal-PFC dynamics are a focus of current theories of mismatch detection. From one perspective, PFC is theorized to play a key role in conveying goal information to support *selective* encoding of novel information that is goal-relevant^70^. Another model posits that predictions and plans for upcoming decisions are generated by the dorsolateral PFC (dlPFC) and relayed to the hippocampus, which later sends mismatch signals to medial PFC^78^. The latter flow of information to medial PFC, depending on whether the hippocampus is conveying a match or a mismatch in expectations, can further bolster the planned action or enable action switching^78^. The dynamics of this information flow imply that control signals mediated by medial PFC are evoked following hippocampal theta mismatch signals, rather than with vmPFC theta driving hippocampal theta^30^. Whether and when PFC subregions drive hippocampal activity, and vice versa, remain unclear. Some studies of hippocampal-PFC dynamics during retrieval have shown that signals from PFC initiate memory retrieval processes in the hippocampus/temporal lobe^79,80^, whereas others demonstrate that hippocampal theta may precede PFC theta during free recall^81^. Other studies of cross-regional communication suggest that information flows from hippocampus to PFC via theta oscillations in both humans^81^ and non-human animals^82,83^. With respect to error-driven dynamics, cross-regional communication following MPEs may reflect engagement in memory encoding as opposed to memory retrieval, and these two processes may exhibit differential hippocampal-PFC dynamics. Altogether, these observations highlight critical open questions about (a) the relationship between MPE-driven PFC and hippocampal activity, (b) the importance of theta oscillations^77^ in mediating cross-regional communication, (c) whether hippocampal-PFC dynamics differ during memory encoding vs. retrieval, and (d) whether the temporal dynamics of hippocampal-PFC information flow vary as a function of PFC subregion. Answers to these questions will advance understanding of cross-regional communication upon MPE detection^70^.

Given limitations in the spatial resolution of scalp EEG, key questions about other components of models of mismatch detection were not examined in the current study. Future work should further examine the role of the nucleus accumbens, which is theorized to integrate goal information from PFC and MPE signals from the hippocampus and the ventral tegmental area/substantia nigra (VTA/SN)^84^. The VTA/SN are posited to release dopamine in response to MPEs to tag hippocampal memory traces for consolidation^70^. Whether dopamine and norepinephrine differentially contribute to learning from MPEs remains unclear. Prior work examining the integrity of LC and SN among older adults implicates the former in episodic memory performance and the latter in working memory^43^. The relative roles of noradrenaline release from LC and dopamine release from VTA/SN^69^ during MPEs warrant further investigation. Notably, the hippocampus receives more profuse projections from LC than VTA neurons in mice^42^. Moreover, LC neurons influence PFC neuronal firing^85,86^ and stimulation of the LC releases both noradrenaline and dopamine in PFC^87,88^. PFC neurons also project to LC^89,90^ and PFC neurons can modulate LC activity^39^. The bidirectional connectivity between LC and PFC raises questions about the temporal dynamics of their interactions following MPEs and how LC modulates PFC-hippocampus interactions.

### Future oriented: How MPEs facilitate future behavior

MPEs can facilitate future behavior by increasing the probability that unexpected information is encoded into memory, thereby allowing internal models of the world to be updated and subsequently improve the accuracy of future predictions^69^. In the current study, the likelihood of learning from strong MPEs was associated with frontal theta magnitude during MPEs. Subsequent memory effects in frontal theta for strong mismatch probes were not selectively observed *during* MPEs, but also prior to and following MPEs; frontal theta during these periods also supported later memory for weak mismatch probes. These findings are consistent with prior work showing that higher peri-stimulus frontal theta boosts memory formation^37,44,45^. Moreover, these findings complement previous research showing that explicit predictions are required for PE-driven learning to take place^11^. We show that memory for expectation-violating information, regardless of the overall strength of the prediction, depended on the amount of control engaged during prediction generation. Although frontal theta *during* MPEs was more strongly associated with memory for strong vs. weak mismatch probes, it did not additionally boost memory for strong mismatch probes above that of weak mismatch probes. A richer assay of memory vividness or precision may reveal differential learning effects as a function of prediction strength. Frontal theta increases elicited at the time of the MPE may reflect PFC conveying the goal relevance of surprising information^70^, hence greater frontal theta in response to strong vs. weak MPEs. Long-term learning resulting from higher frontal theta in response to strong MPEs may be mediated by hippocampal learning mechanisms and therefore may be further enhanced by consolidation. Given that scalp EEG was not acquired in Experiment 2 (recognition memory test 48hrs later), it remains unclear whether the subsequent memory effects in frontal theta hold after allowing for memory consolidation. Whether control-associated enhancements in learning are specific to learning from MPEs can be further tested in future work by including a test of subsequent memory for weak match probes as an additional control condition for comparison.

We hypothesized that strong MPE-elicited pupillary size increases, indexed by PC3 and PC4, would support new learning, given theories of LC-hippocampus interactions, the impact of these interactions on memory consolidation^19^, prior findings of pupil-linked, PE-related learning effects^11^, and prior findings that unsigned reward PEs – presumably signaled by the LC-norepinephrine system – enhance memory^91^. To our surprise, neither PC3 nor PC4 scores were associated with subsequent recognition memory for mismatch items on the same day (Experiment 1) or after a longer delay that allowed for overnight consolidation (48hrs later; Experiment 2). Whether a more hippocampal-dependent subsequent memory assay would reveal pupil-related and consolidation-dependent learning effects remains an open question for future research.

Finally, the magnitude of observed increases in control, attention, and arousal following strong MPEs may be influenced by the decision-making process engaged when making match/mismatch judgments. Not all MPEs necessitate behavioral responses. Whether similar magnitudes in neurocognitive responses and consequences for learning are observed upon detection of an MPE, but in the absence of a decision remains unclear.

Mnemonic predictions guide everyday behavior. The strength of these predictions varies depending on frequency of prior occurrences^21^, emotional salience^92,93^, and attention during learning^41,94,95^, amongst other factors. Here, we report that violations of *strong* mnemonic predictions elicited increases in cognitive control, attention, and arousal. Furthermore, the magnitude of cognitive control responses to strong MPEs modulated the likelihood of remembering that information in the future. Altogether, these findings advance understanding of the learning and retrieval rules of the mind and brain and add to a growing literature on how dynamic interactions between knowledge about the past and the present give rise to rapid shifts in cognition that impact learning and future remembering.

## Acknowledgments

This work was supported by the National Institute on Aging R01AG065255 to A.D.W., National Science Foundation GRF DGE-2146755 to A.M.X., and a Stanford Interdisciplinary Graduate Fellowship to A.M.X. The authors would like to thank the members of the Stanford Memory Lab for useful discussions.

## Footnotes

a n.b., At the trial-level in Fig. 2e, there was no Mismatch × frontal theta interaction on posterior alpha and no main effect of frontal theta on posterior alpha; however, the posterior alpha mediation model (Fig. 3c), wherein the predictors were additionally rescaled, suggests that frontal theta and posterior alpha may be negatively associated at a trial-level following a strong MPE.

## Notes

### Competing Interest Statement

The authors have declared no competing interest.

### Summary of Updates

Sections 2 and 3 were removed to help make key takeaways clear. Summary figure updated.

