## Supplemental Figures and Tables for "Strong mnemonic prediction errors increase cognitive control, attention, and arousal"

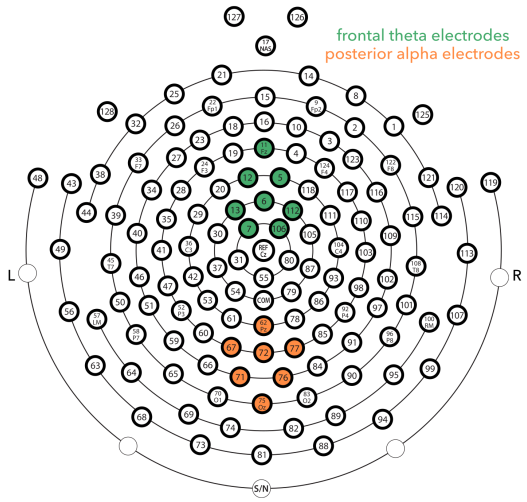


**Supplementary Fig. 1. Frontal theta and posterior alpha electrodes.** Depiction of electrodes analyzed to compute frontal theta (green) and posterior alpha power (orange).

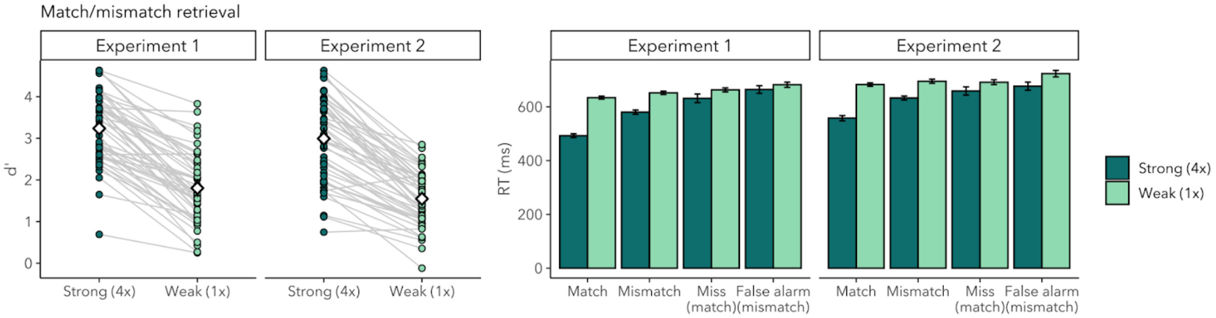


**Supplementary Fig. 2. Behavior in the associative retrieval task in Experiment 1 and Experiment 2.** Associative memory *d'* for strong pairings was 3.24±0.83 in Experiment 1 and 2.99±1.00 in Experiment 2. Associative memory *d'* for weak pairings was 1.80±0.84 in Experiment 1 and 1.55±0.65 in Experiment 2.

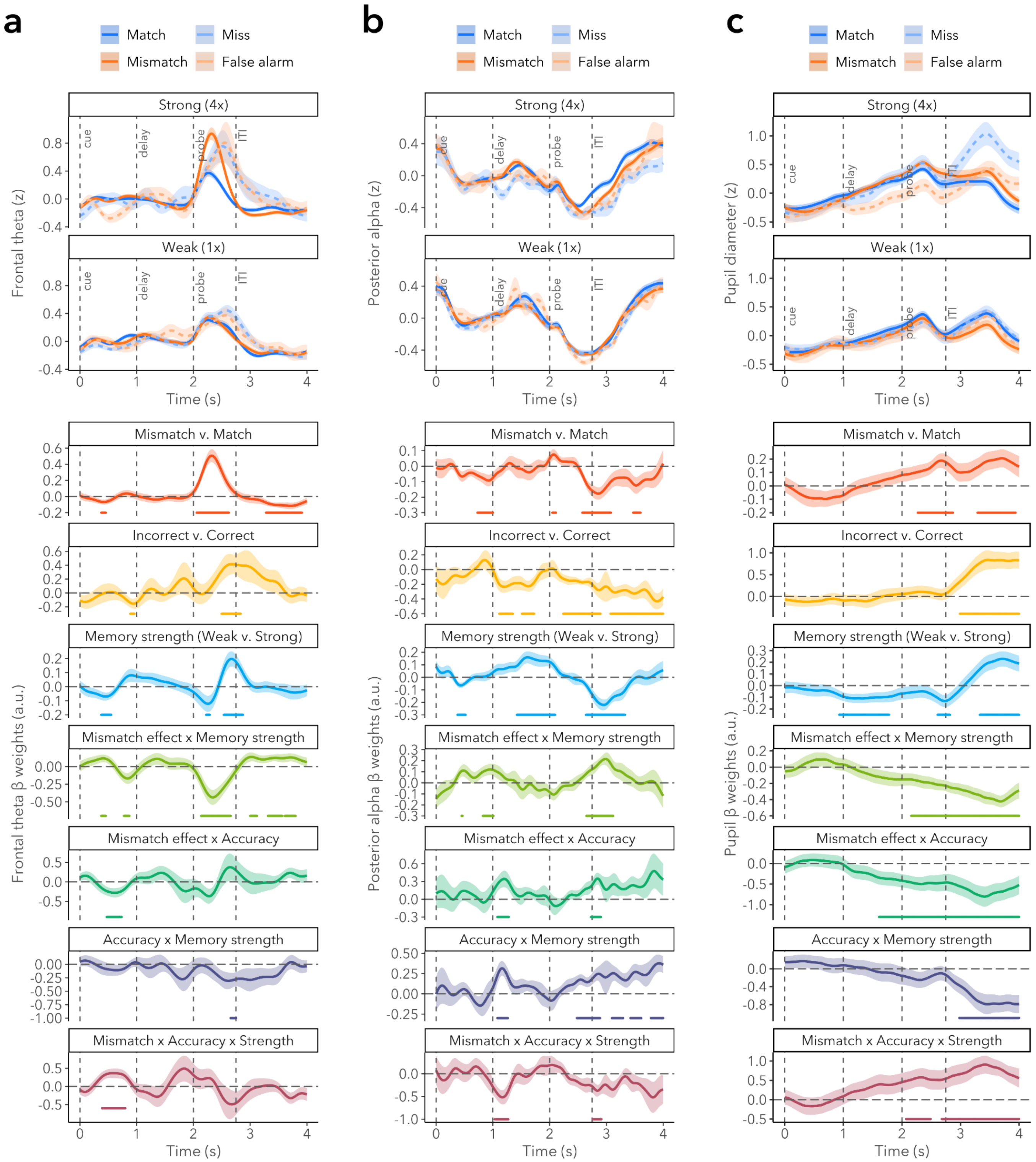


**Supplementary Fig. 3. Trial-level regression plots including miss and false alarm trials.** (a) Clusters for β weights for frontal theta ~ Mismatch * Accuracy *Strength across 25 subjects. (b) Clusters for β weights for posterior alpha ~ Mismatch * Accuracy *Strength across 25 subjects. Near probe onset, there was additionally a main effect of Mismatch (1.976 to 2.128s; β=0.06, CI=[0.02, 0.10]; p<0.001) and a Mismatch × Strength interaction (1.952 to 2.112s; β=-0.09, CI=[-0.15, -0.02]; p<0.001). These effects may reflect the offset of the delay period Strength effect or anticipation of the oncoming probe. (c) Clusters for β weights for pupil diameter ~ Mismatch * Accuracy * Strength across 31 subjects. For statistics on clusters, see Supplementary Table 3.

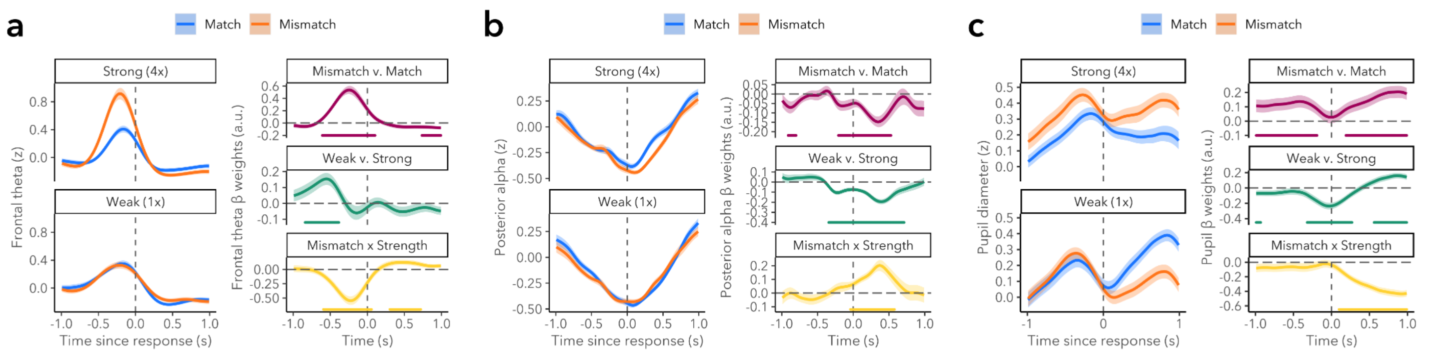


**Supplementary Fig. 4. Response-locked trial-level regression plots.** (a) Response-locked frontal theta analyses showed that before responses, there was a main effect of Mismatch (-0.608 to 0.104s; β=0.33, CI=[0.24, 0.42]; p<0.001), a main effect of Strength (-0.840 to -0.384s; β=0.12, CI=[0.05, 0.19]; p<0.001), and a Mismatch × Strength interaction (-0.592 to 0.056s; β=-0.36, CI=[-0.47 -0.25]; p<0.001). The interaction indicated that the difference in frontal theta between mismatch and match trials was larger for strong compared to weak trials. After responses, there was a main effect of Mismatch (0.736 to 0.992s; β=-0.07, CI=[-0.12, -0.02]; p<0.001) that emerged later than a Mismatch × Strength interaction (0.304 to 0.720s; β=0.11, CI=[0.04, 0.19]; p<0.001). Together, these results indicate that after responses, the differences in the magnitude of the dip in frontal theta between mismatch and match trials was larger for weak compared to strong trials; differences between mismatch-evoked decreases in frontal theta persisted. (b) Response-locked posterior alpha analyses showed that responses coincided with a drop in posterior alpha power. There were main effects of Mismatch (-0.208 to -0.040s; β=-0.06, CI=[-0.109, -0.011]; p<0.001; and from -0.016 to 0.528s; β=-0.10, CI=[-0.14, -0.06]; p<0.001), a main effect of Strength (-0.344 to 0.712s; β=-0.12, CI=[-0.15, -0.09]; p<0.001), and a Strength × Mismatch interaction (-0.040 to 0.576s; β=0.14, CI=[0.09, 0.18]; p<0.001). The difference in posterior alpha power between match and mismatch trials was therefore larger for strong compared to weak trials. The onset of the interaction was later than that of the Mismatch and Strength effects, indicating strength-modulated changes in attention following MPEs emerged primarily after responses. There was additionally a Mismatch effect from -0.912 to -0.800s (β=-0.07, CI=[-0.12, -0.01]). (c) Response-locked analyses of changes in pupil diameter revealed that there were main effects of Mismatch (-0.99 to -0.19; β=0.12, CI=[0.04, 0.20], p<0.001; 0.19 to 0.99s; β=0.16, CI=[0.08, 0.24]; p<0.001), main effects of Strength (-0.99 to -0.93s; β=-0.07, CI=[-0.14, 0]; p<0.001; -0.32 to 0.27, β=-0.17, CI=[-0.25, -0.10], p<0.001; 0.56 to 0.99s, β=0.14, CI=[0.06, 0.21], p<0.001), and a Mismatch × Strength interaction (0.10 to 0.99s; β=-0.33, CI=[-0.41, -0.25]; p<0.001). The interaction indicates that the difference in pupil size between match and mismatch trials was larger for strong compared to weak trials.

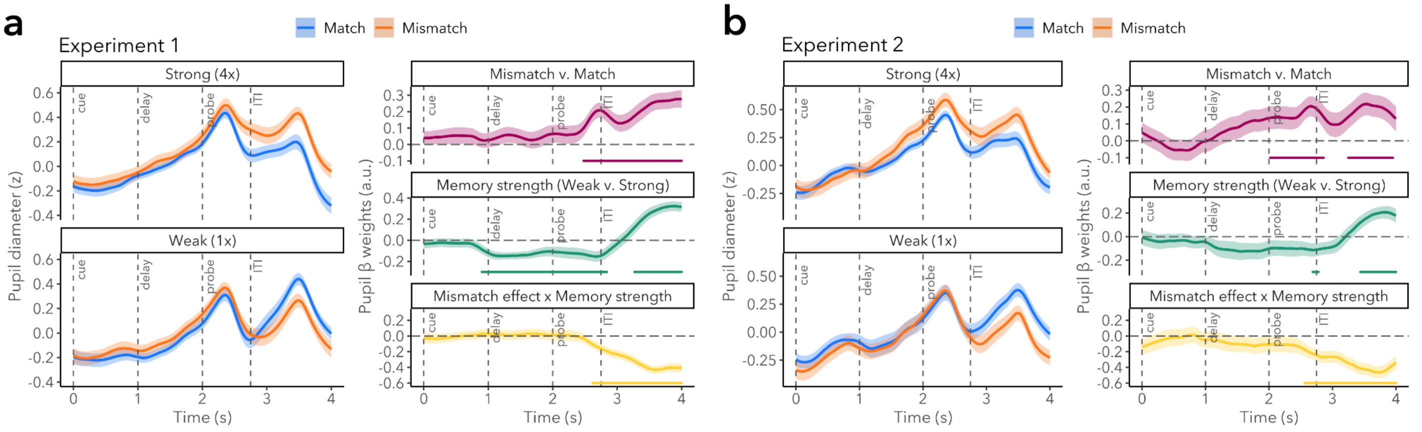


**Supplementary Fig. 5. Pupil trial-level regression plots separated by experiment.** Effects of Mismatch and Strength on pupil diameter in Experiment 1 (a) and Experiment 2 (b). (a) In Experiment 1, there was a main effect of Mismatch from 2.48 to 4.00s (β=0.20, CI=[0.11, 0.29], p<0.001), main effects of Strength from 0.90 to 2.84s (β=-0.13, CI=[-0.21, -0.05]) and 3.27 to 4.00s (β=0.27, CI=[0.17, 0.37]), and a Mismatch × Strength interaction from 2.62 to 4.00s (β=-0.31, CI=[-0.40, -0.22]). (b) In Experiment 2, there were main effects of Mismatch from 2.02 to 2.86s (β=0.16, CI=[0.04, 0.28], p<0.001) and

3.25 to 3.95s (β=0.19, CI=[0.07, 0.32]), main effects of Strength from 2.69 to 2.78s (β=-0.11, CI=[-0.22, 0]) and 3.44 to 4.00s (β=0.18, CI=[0.08, 0.28]), and a Mismatch × Strength interaction from 2.56 to 4.00s (β=-0.34, CI=[-0.47, -0.21]).

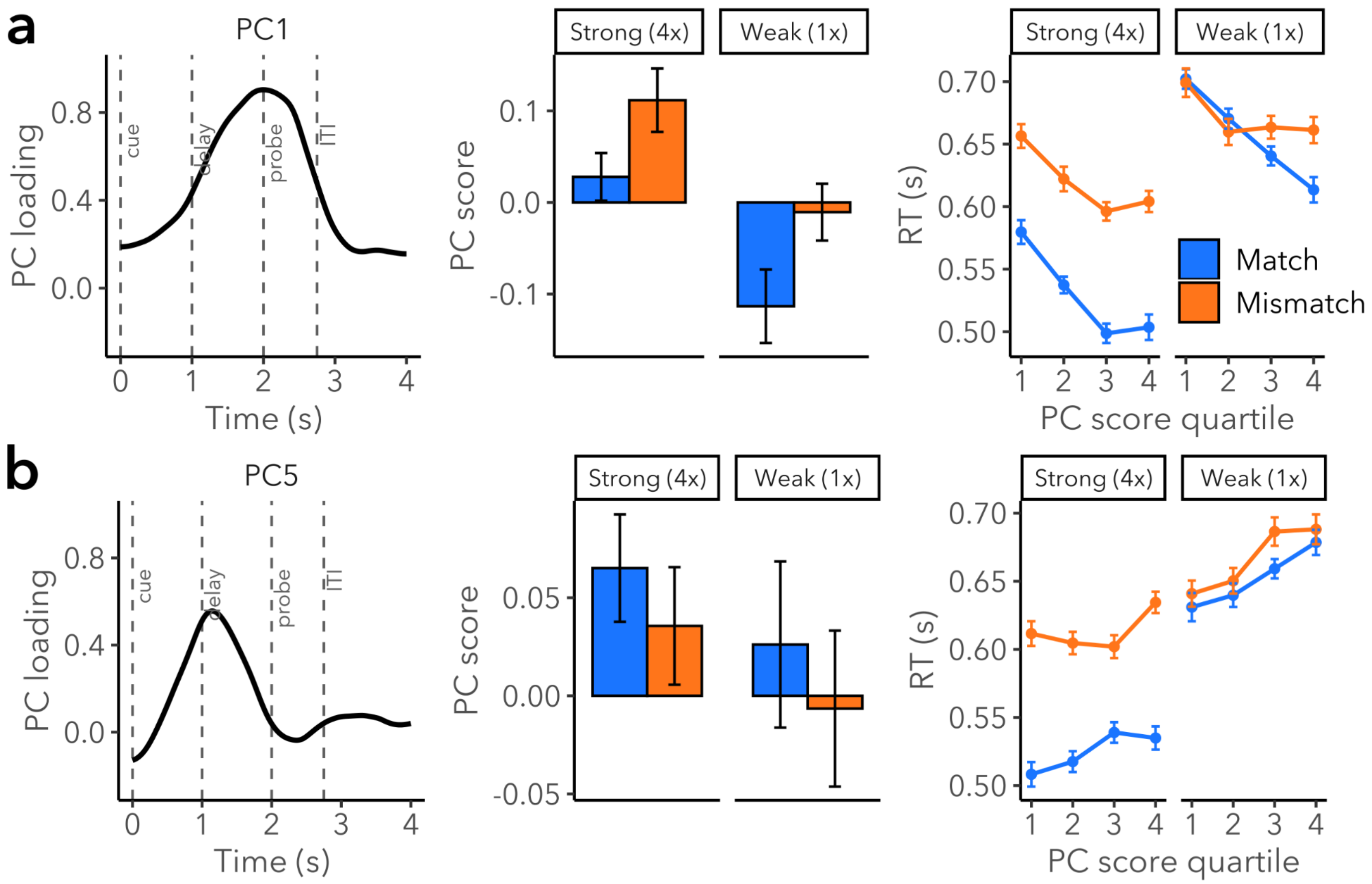


**Supplementary Fig. 6. Retrieval success and cognitive effort (PC1 and PC5) scores.** (a) PC1 scores were higher for stronger memories both at the condition level and the trial level, leading us to interpret PC1 as indicative of the strength of memory retrieval. In particular, PC1 scores were higher for strong compared to weak trials (**Supplementary Table 4**; effect of Strength) and higher PC1 scores predicted faster match RTs (**Supplementary Table 5**; effect of PC1 score). The relationship between PC1 scores and RTs was attenuated for mismatch trials (Mismatch × PC1 score interaction), which may be a consequence of the response delay induced by MPEs on mismatch trials (**Fig. 2c**). PC1 accounted for the most variance (28.81%), onset during the cue period, peaked predominantly at the time of probe onset (2.00s, CI=[1.94, 2.05]; loading = 0.90, CI=[0.90, 0.91]), and had a FWHM of 1.75s (CI=[1.70, 1.78]). Altogether, these findings suggest that PC1 tracks graded retrieval success or strength of memory retrieval. (b) For PC5, the relationship with the memory strength conditions and RTs indicated that this component was a condition-level marker of cue strength and a trial-level marker of cognitive effort during associative retrieval. PC5 scores were higher for strong than weak trials (**Supplementary Table 6**; main effect of Strength), but in contrast to PC1, higher PC5 scores were associated with longer RTs (**Supplementary Table 7**; main effect of PC5 scores). The relationship between PC5 scores and RTs was stronger for weak compared to strong pairings (PC5 score × Strength). Thus, PC5 showed the opposite pattern as PC1, with PC5 scores scaling inversely with trial-level memory strength, suggesting that PC5 reflects a mixture of cue strength (i.e., higher scores for strong cues) and the amount of cognitive effort expended during associative memory retrieval (i.e., inverse relationship with RT). The temporal profile of PC5 is consistent with this interpretation, as PC5 onset during the cue period, peaked during the delay period (1.15s after cue onset, CI=[1.12, 1.18]; loading = 0.56, CI=[0.52, 0.61]) and had a FWHM of 0.98s (CI=[0.92, 1.04]). This component accounted for 5.66% of variance.

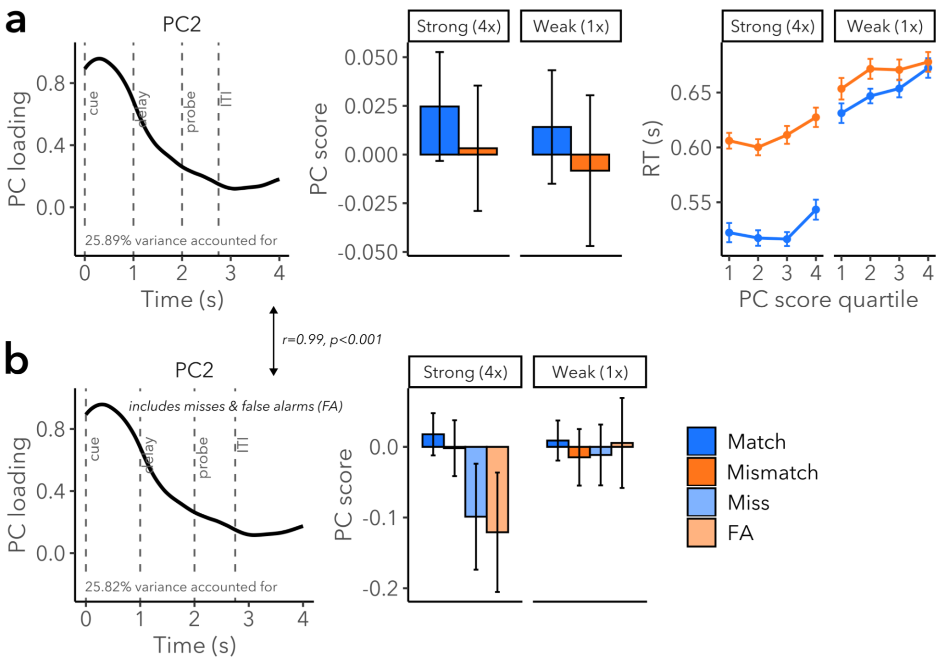


**Supplementary Fig. 7. Comparison of preparatory attention pupil (PC2) scores computed with and without miss and false alarm trials.** (a) PC2 peaked 0.30s (CI=[0.29, 0.31]) after cue onset (loading = 0.96, CI=[0.95, 0.96]) and declined slowly over the course of the trial, explaining 25.89% of variance. There was a marginal difference in PC2 scores as a function of Mismatch, but not as a function of Strength (**Supplementary Table 8**). PC2 scores were marginally associated with longer RTs (**Supplementary Table 9**; marginal main effect of PC2 scores). (b) An augmented model using scores computed in a tPCA including error trials revealed a main effect of Accuracy and an Accuracy × Strength interaction (**Supplementary Table 10**). The augmented model did not fit the data significantly better than a compact model lacking Accuracy terms ($\chi2(4)=7.66$, $p=0.105$). Together, these outcomes indicate that the influence of pupil-linked arousal immediately following cue presentation on memory retrieval influences the likelihood of successful memory retrieval and may impact the precision with which a memory is retrieved.

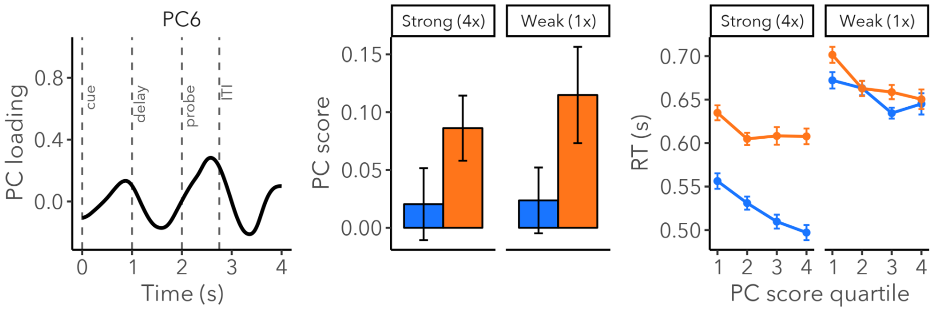


**Supplementary Fig. 8. Stimulus detection and mismatch detection (PC6) scores.** PC6 explained the least amount of variance (1.83%) and had its most prominent peak during probe presentation (0.58s after probe onset, CI=[0.55, 0.60]; loading = 0.28, CI=[0.26, 0.30]; FWHM=0.60, CI=[0.56, 0.64]); the second most prominent peak occurred earlier, 0.87s after cue onset (CI=[0.85, 0.90]; loading = 0.13, CI=[0.11, 0.16]; FWHM=0.45, CI=[0.43, 0.47]); the last peak was at the end of the trial (3.98s after cue onset, CI=[3.94, 3.99]; loading = 0.10, CI=[0.08, 0.11]). The temporal profile of PC6's loadings suggest that this component reflects responses to changes in luminance driven by stimulus onset; these responses peak after ~500-900ms. Given that PC6's loadings were higher for the probe-period response compared to the cue-period response, PC6 may be more sensitive to the identity of the probe than that of the cue. PC6 scores were higher on mismatch than match trials (**Supplementary Table 11**; main effect of Mismatch) and higher PC6 scores predicted faster RTs (**Supplementary Table 12**; main effect of PC6 score), indicating that PC6 scores may reflect the amount of attention allocated to the probe and its identity as a match or mismatch. There was additionally a PC6 score × Mismatch interaction, showing that the relationship with RTs was attenuated for mismatches. Unlike PC3 and PC4, PC6 scores (from the full dataset with error trials) were not fit better by an augmented model including effects of and interactions with Accuracy

($\chi^2(4)=4.31$, $p=0.366$). Misses and false alarms can be accounted for by an erroneous mnemonic prediction (which could give rise to a MPE) or the absence of a prediction (where the response may reflect guessing). Given ambiguity regarding the experience of MPEs on error trials, and PC6's lack of sensitivity to accuracy, PC6 may not be fully characterized as an MPE-sensitive component, but rather an initial mismatch response.

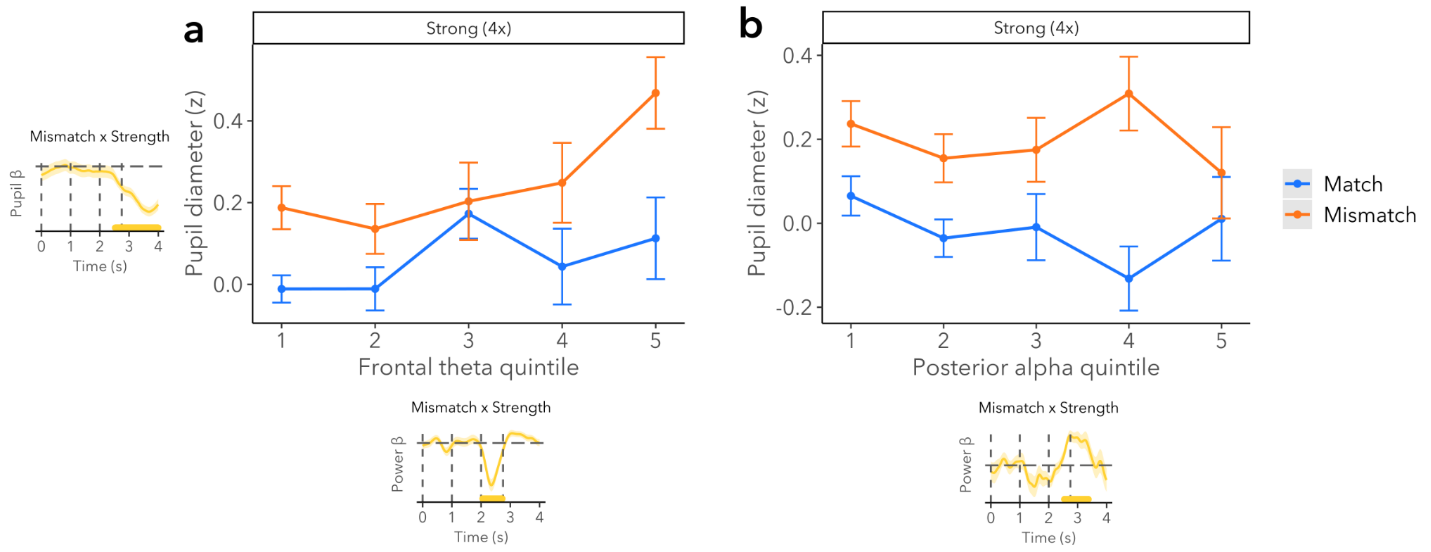


**Supplementary Fig. 9. Association of pupillary MPE-related responses to frontal theta and posterior alpha.** Mean pupil responses during strong MPEs are related to frontal theta and posterior alpha responses during strong MPEs, which did not differ with Mismatch. (a) Relationship between frontal theta and mean pupil. (b) Relationship between posterior alpha and mean pupil. Statistics are reported in the main text.

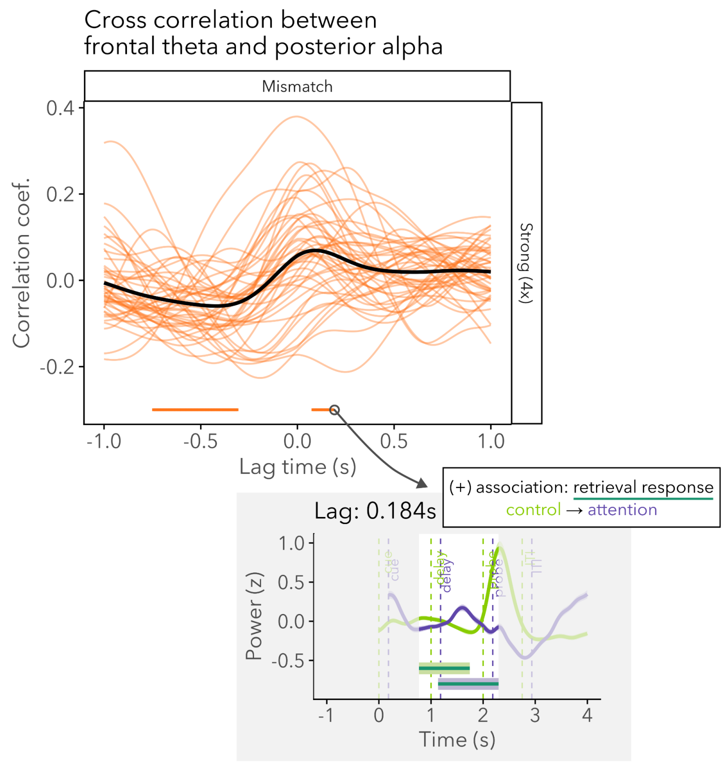


**Supplementary Fig. 10. Cross-correlation of frontal theta and posterior alpha reveals temporal relationship of retrieval-related processes.** The cross-correlation figure is a reproduction of that in **Fig. 4d**; the inset in gray shows the mean frontal theta and posterior alpha time series from **Fig. 2d** and **Fig. 2e** with posterior alpha lagging at the max timepoint in the indicated cross-correlation cluster (0.184s). This cross-correlation analysis of the frontal theta and posterior alpha time series revealed a significant cluster (a) with a positive correlation and (b) with frontal theta leading (**Fig. 4d**; 0.080 to 0.184s, r=0.006, CI=[0.065, 0.068]). This positive association – greater control and less attention – may reflect cognition during memory retrieval, given the positive mean β during the delay-period Strength clusters in **Fig. 2d** and **Fig 2e**. The positive lag suggests that at retrieval, cognitive control is first engaged and changes in attention, perhaps toward retrieved contents, follows.

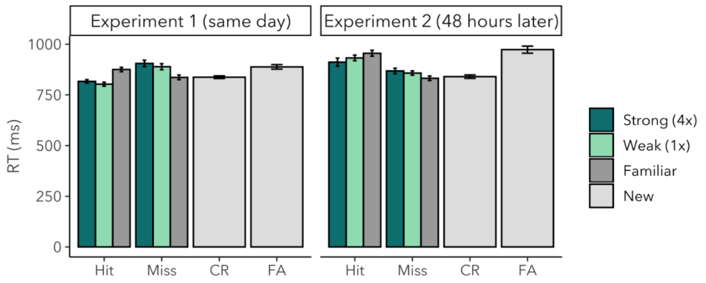


**Supplementary Fig. 11. Subsequent memory RTs for mismatch probes in Experiments 1 and 2.** A linear mixed effects model was used to examine how recognition RTs in the subsequent memory test varied with Strength, Accuracy, Experiment, and their interactions. Pairwise comparisons revealed that correct RTs for strong probes were faster than for familiar probes in the same day test ($\Delta_{Strong-Familiar}$=-0.050, CI=[-0.079, -0.021], p<0.001) but not 48 hrs later ($\Delta_{Strong-Familiar}$=-0.019, CI=[-0.056, 0.019], p=0.583). Correct RTs for strong probes were faster than for new probes in the same day test ($\Delta_{Strong-New}$=-0.036, CI=[-0.056, -0.015], p<0.001) and 48 hrs later ($\Delta_{Strong-New}$=0.061, CI=[0.033, 0.089], p<0.001). Correct RTs between strong and weak probes did not differ in the same day test ($\Delta_{Strong-Weak}$=0.012, CI=[-0.013, 0.038], p=0.592) or 48 hrs later ($\Delta_{Strong-Weak}$=-0.014, CI=[-0.050, 0.023], p=0.769). These results reveal that an MPE-related benefit on subsequent memory RTs diminished when recognition memory was tested 48 hrs later. Altogether, subsequent recognition performance revealed that memory was superior for probes that violated strong or weak mnemonic predictions than for familiar items that were not encountered in the context of an MPE. When memory is tested after a 48-hr delay, these effects diminish.

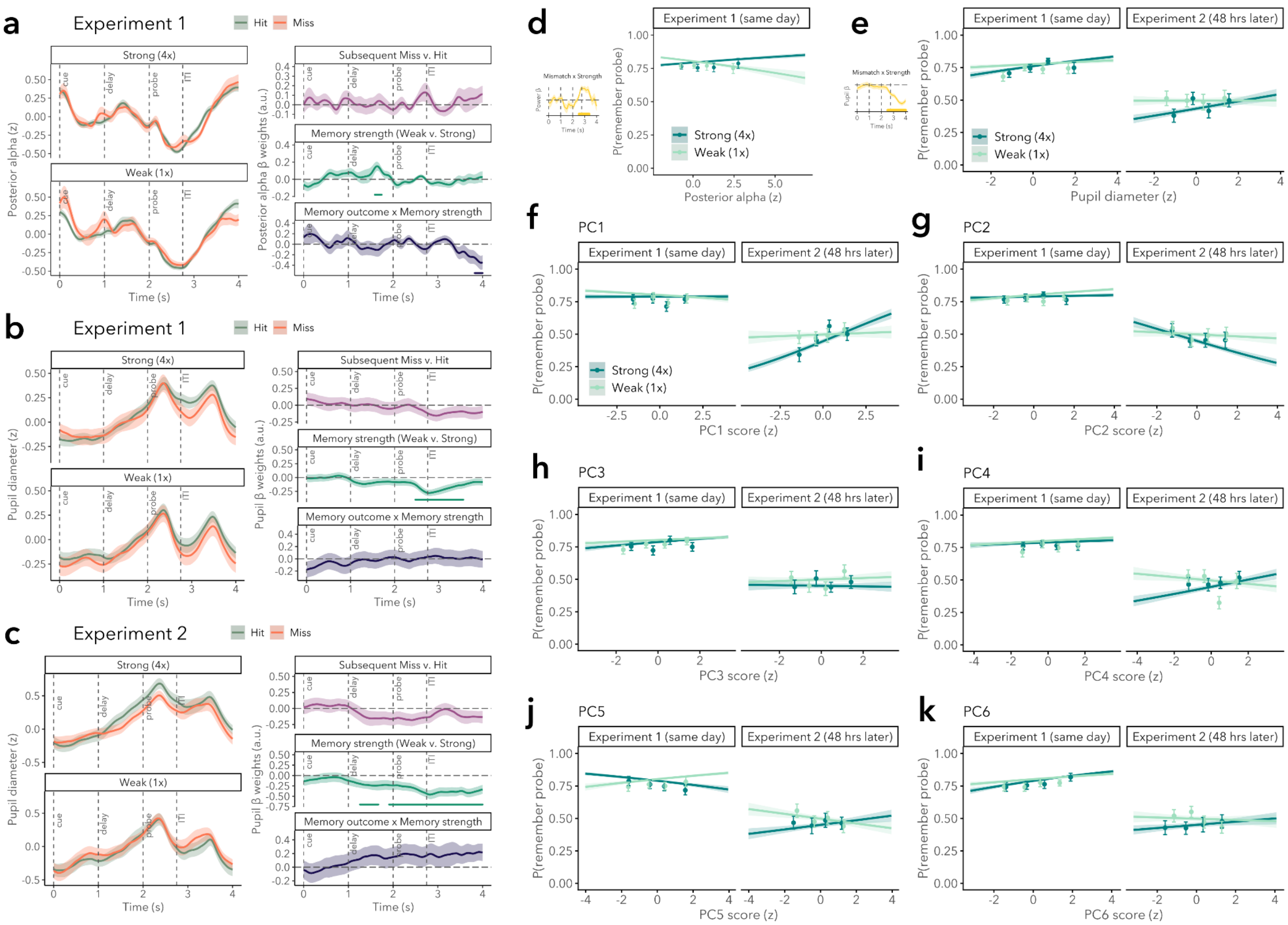


**Supplementary Fig. 12. Neural substrates underlying learning from MPEs.** (a) Trial-level regression models revealed that for posterior alpha, there was a memory outcome × Strength interaction from 1.824–2.000s (β=-0.32, CI=[-0.56, -0.08]) after probe onset such that the difference in posterior alpha power between hits and misses was larger for weak compared to strong trials. There was a main effect of Strength from 1.592 to 1.744s after cue onset (β=0.134, CI=[0.039, 0.228]). (b–c) For pupil size, there were no windows of time in Experiment 1 or Experiment 2 which predicted subsequent memory. (b) In Experiment 1, there was a main effect of Strength from 0.47 to 1.56s after probe onset (β=-0.21, CI=[-0.32, -0.09]). (c) In Experiment 2, there were main effects of Strength from 1.26 to 1.67s (β=-0.23, CI=[-0.42, -0.04]) and 1.91 to 4.00s (β=-0.36, CI=[-0.54, -0.18]). (d–e) Subsequent memory effects for mean posterior alpha power and mean pupil diameter during the primary Mismatch × Strength clusters, illustrated by the yellow clusters, were not found. (d) There was no effect of subsequent memory outcome on mean MPE-related posterior alpha (main effect of subsequent memory outcome: β=-0.030, CI=[-0.089, 0.030], p=0.326; memory outcome × Strength: β=0.038, CI=[-0.047, 0.123], p=0.377. (e) There was no effect of subsequent memory outcome on mean MPE-related pupil size (main effect of subsequent memory outcome: β=-0.085, CI=[-0.216, 0.047], p=0.205; memory outcome × Strength: β=0.036, CI=[-0.152, 0.225], p=0.705; memory outcome × experiment: β=-0.034, CI=[-0.228, 0.159], p=0.728; no three-way interaction: β=0.113, CI=[-0.164, 0.390], p=0.425). (f) PC1 scores did not predict subsequent memory (no main effect of PC1: O.R.=1.000, CI=[0.852, 1.174], p=0.995; no PC1 × Strength interaction: O.R=0.950, CI=[0.753, 1.198], p=0.753; a marginal PC1 × Experiment interaction: O.R.=1.246, CI=[0.977, 1.589], p=0.077; no PC1 × Strength × Experiment interaction: O.R.=0.862, CI=[0.609, 1.219], p=0.400). (g) PC2 scores did not predict subsequent memory (no main effect of PC2: O.R.=1.018, CI=[0.868, 1.193], p=0.826; no PC2 × Strength interaction: O.R.=1.061, CI=[0.841, 1.338], p=0.619; a marginal PC2 × Experiment interaction: O.R.=0.814, CI=[0.641, 1.035], p=0.093; no PC2 × Strength × Experiment interaction: O.R.=1.103, CI=[0.781, 1.559], p=0.578). (h) PC3 scores did not predict subsequent memory (no main effect of PC3: O.R.=1.079, CI=[0.921, 1.264], p=0.348; no PC3 × Strength interaction: O.R.=0.968, CI=[0.766, 1.225], p=0.789; no PC3 × Experiment interaction: O.R.=0.918, CI=[0.723, 1.166], p=0.485; no PC3 × Strength × Experiment interaction: O.R.=1.067, CI=[0.752, 1.513], p=0.717). (i) PC4 scores did not predict subsequent memory (no main effect of PC4: O.R.=1.030, CI=[0.879, 1.207], p=0.716; no PC4 × Strength interaction: O.R.=1.018, CI=[0.806, 1.284], p=0.883; no PC4 × Experiment interaction: O.R.=1.086, CI=[0.858, 1.375], p=0.493; no PC4 × Strength × Experiment interaction: O.R.=0.834, CI=[0.589, 1.181], p=0.307). (j) PC5 scores did not predict subsequent memory (no main effect of PC5: O.R.=0.912, CI=[0.782, 1.064], p=0.240; no PC5 × Strength interaction: O.R.=1.195, CI=[0.947, 1.507], p=0.133; no PC5 × Experiment interaction: O.R.=1.178, CI=[0.935, 1.485], p=0.165; marginal PC5 × Strength × Experiment interaction: O.R.=0.723, CI=[0.512, 1.022], p=0.067). (k) PC6 scores did not predict subsequent memory (marginal main effect of PC6: O.R.=1.138, CI=[0.979, 1.323], p=0.093; no PC6 × Strength interaction: O.R.=0.944, CI=[0.746, 1.194], p=0.631; no PC6 × Experiment interaction: O.R.=0.924, CI=[0.737, 1.158], p=0.490; no PC6 × Strength × Experiment interaction: O.R.=0.989, CI=[0.697, 1.403], p=0.951).

**Supplementary Table 1: Frontal theta trial-level regression model including miss and false alarm trials**

| | Start | End | Mean β [95% CI] |
|---|---|---|---|
| Mismatch v. Match | 2.064 | 2.640 | 0.34 [0.23, 0.45] |
| Mismatch v. Match | 3.296 | 3.928 | -0.10 [-0.15, -0.05] |
| Incorrect v. Correct | 2.536 | 2.888 | 0.55 [0.16, 0.93] |
| Memory strength (Weak v. Strong) | 0.360 | 0.528 | -0.06 [-0.12, -0.01] |
| Memory strength (Weak v. Strong) | 2.520 | 2.880 | 0.17 [0.08, 0.27] |
| Mismatch effect x Memory strength | 0.776 | 0.848 | -0.15 [-0.30, -0.01] |
| Mismatch effect x Memory strength | 2.136 | 2.672 | -0.35 [-0.51, -0.18] |
| Mismatch effect x Memory strength | 3.032 | 3.096 | 0.12 [0.01, 0.24] |
| Mismatch effect x Memory strength | 3.384 | 3.488 | 0.11 [0.01, 0.20] |
| Mismatch effect x Memory strength | 3.640 | 3.816 | 0.12 [0.02, 0.23] |
| Mismatch effect x Accuracy | 0.536 | 0.776 | -0.26 [-0.47, -0.06] |
| Mismatch effect x Accuracy | 2.232 | 2.312 | -0.50 [-0.96, -0.05] |
| Accuracy x Memory strength | 2.640 | 2.808 | -0.41 [-0.76, -0.06] |
| Mismatch x Accuracy x Strength | 0.424 | 0.824 | 0.35 [0.15, 0.55] |
| Mismatch x Accuracy x Strength | 3.648 | 3.792 | -0.37 [-0.68, -0.05] |

**Supplementary Table 2: Posterior alpha trial-level regression model including miss and false alarm trials**

| | Start | End | Mean β [95% CI] |
|---|---|---|---|
| Mismatch v. Match | 0.760 | 1.000 | -0.08 [-0.14, -0.02] |
| Mismatch v. Match | 2.040 | 2.104 | 0.07 [0.01, 0.13] |
| Mismatch v. Match | 2.576 | 3.064 | -0.14 [-0.20, -0.07] |
| Mismatch v. Match | 3.448 | 3.648 | -0.12 [-0.21, -0.02] |
| Incorrect v. Correct | 1.096 | 1.376 | -0.22 [-0.37, -0.08] |
| Incorrect v. Correct | 1.504 | 1.736 | -0.21 [-0.34, -0.07] |
| Incorrect v. Correct | 2.240 | 2.904 | -0.18 [-0.26, -0.09] |

| | | | |
|---|---|---|---|
| Incorrect v. Correct | 3.072 | 4.000 | -0.31 [-0.43, -0.19] |
| Memory strength (Weak v. Strong) | 0.376 | 0.520 | -0.06 [-0.10, -0.02] |
| Memory strength (Weak v. Strong) | 1.312 | 2.096 | 0.13 [0.06, 0.20] |
| Memory strength (Weak v. Strong) | 2.640 | 3.352 | -0.16 [-0.22, -0.09] |
| Mismatch effect x Memory strength | 0.840 | 1.016 | 0.11 [0.02, 0.20] |
| Mismatch effect x Memory strength | 2.648 | 3.128 | 0.16 [0.09, 0.24] |
| Mismatch effect x Accuracy | 1.088 | 1.280 | 0.30 [0.06, 0.54] |
| Mismatch effect x Accuracy | 2.736 | 2.904 | 0.29 [0.11, 0.46] |
| Accuracy x Memory strength | 1.072 | 1.272 | 0.28 [0.12, 0.43] |
| Accuracy x Memory strength | 2.488 | 2.896 | 0.18 [0.06, 0.29] |
| Accuracy x Memory strength | 3.096 | 3.224 | 0.22 [0.05, 0.39] |
| Accuracy x Memory strength | 3.440 | 3.640 | 0.25 [0.08, 0.43] |
| Accuracy x Memory strength | 3.752 | 4.000 | 0.34 [0.14, 0.54] |
| Mismatch x Accuracy x Strength | 1.032 | 1.240 | -0.42 [-0.70, -0.15] |
| Mismatch x Accuracy x Strength | 2.768 | 2.912 | -0.32 [-0.56, -0.09] |

**Supplementary Table 3: Pupil trial-level regression model including miss and false alarm trials**

| | Start | End | Mean β [95% CI] |
|---|---|---|---|
| Mismatch v. Match | 2.27 | 2.87 | 0.15 [0.05, 0.25] |
| Mismatch v. Match | 3.29 | 3.94 | 0.18 [0.06, 0.31] |
| Incorrect v. Correct | 2.99 | 4.00 | 0.74 [0.37, 1.11] |
| Memory strength (Weak v. Strong) | 0.93 | 1.78 | -0.10 [-0.18, -0.03] |
| Memory strength (Weak v. Strong) | 2.61 | 2.82 | -0.12 [-0.22, -0.02] |
| Memory strength (Weak v. Strong) | 3.33 | 4.00 | 0.20 [0.08, 0.33] |
| Mismatch effect x Memory strength | 2.16 | 4.00 | -0.29 [-0.41, -0.17] |
| Mismatch effect x Accuracy | 1.61 | 4.00 | -0.55 [-0.91, -0.19] |

| | | | |
|---|---|---|---|
| Accuracy x Memory strength | 2.98 | 4.00 | -0.70 [-1.07, -0.33] |
| Mismatch x Accuracy x Strength | 2.07 | 2.49 | 0.54 [0.08, 1.00] |
| Mismatch x Accuracy x Strength | 2.68 | 4.00 | 0.72 [0.31, 1.13] |

**Supplementary Table 4: PC1 scores as a function of Mismatch and Strength**

| Fixed effects | Estimate | CI | p-value | SE | DF |
|---|---|---|---|---|---|
| (Intercept) | 0.038 | [-0.036, 0.113] | 0.312 | 0.038 | 112.302 |
| Mismatch | 0.032 | [-0.021, 0.085] | 0.242 | 0.027 | 10269.199 |
| Strength | -0.102 | [-0.147, -0.057] | 0.000 | 0.023 | 10271.457 |
| Mismatch x Strength | 0.027 | [-0.052, 0.106] | 0.506 | 0.040 | 10264.248 |

**Supplementary Table 5: PC1 scores and RTs**

| Fixed effects | Estimate | CI | p-value | SE | DF |
|---|---|---|---|---|---|
| (Intercept) | -0.655 | [-0.859, -0.452] | 0.012 | 0.025 | 1.234 |
| PC1 score | -0.050 | [-0.059, -0.040] | 0.000 | 0.005 | 170.709 |
| Strength | 0.201 | [0.185, 0.216] | 0.000 | 0.008 | 104.316 |
| Mismatch | 0.159 | [0.142, 0.176] | 0.000 | 0.009 | 136.383 |
| PC1 score x Strength | -0.004 | [-0.014, 0.007] | 0.479 | 0.005 | 9151.269 |
| PC1 score x Mismatch | 0.021 | [0.008, 0.034] | 0.001 | 0.006 | 9112.367 |
| Strength x Mismatch | -0.122 | [-0.141, -0.104] | 0.000 | 0.009 | 10189.081 |
| PC1 score x Strength x Mismatch | 0.014 | [-0.004, 0.033] | 0.123 | 0.009 | 10218.503 |

**Supplementary Table 6: PC5 scores as a function of Mismatch and Strength**

| Fixed effects | Estimate | CI | p-value | SE | DF |
|---|---|---|---|---|---|

| | | | | | |
|---|---|---|---|---|---|
| (Intercept) | 0.081 | [-0.353, 0.515] | 0.301 | 0.045 | 1.137 |
| Mismatch | -0.041 | [-0.094, 0.012] | 0.127 | 0.027 | 10258.839 |
| Strength | -0.090 | [-0.134, -0.046] | 0.000 | 0.023 | 10260.729 |
| Mismatch x Strength | 0.045 | [-0.033, 0.124] | 0.257 | 0.040 | 10255.146 |

**Supplementary Table 7: PC5 scores and RTs**

| Fixed effects | Estimate | CI | p-value | SE | DF |
|---|---|---|---|---|---|
| (Intercept) | -0.663 | [-1.107, -0.219] | 0.033 | 0.036 | 1.016 |
| PC5 score | 0.015 | [0.007, 0.022] | 0.000 | 0.004 | 221.355 |
| Strength | 0.215 | [0.205, 0.226] | 0.000 | 0.005 | 10252.177 |
| Mismatch | 0.163 | [0.151, 0.176] | 0.000 | 0.006 | 10232.294 |
| PC5 score x Strength | 0.010 | [-0.001, 0.020] | 0.066 | 0.005 | 9762.962 |
| PC5 score x Mismatch | -0.008 | [-0.021, 0.004] | 0.203 | 0.006 | 10076.592 |
| Strength x Mismatch | -0.127 | [-0.145, -0.108] | 0.000 | 0.009 | 10232.382 |
| PC5 score x Strength x Mismatch | -0.002 | [-0.021, 0.016] | 0.796 | 0.010 | 10158.572 |

**Supplementary Table 8: PC2 scores as a function of Mismatch and Strength**

| Fixed effects | Estimate | CI | p-value | SE | DF |
|---|---|---|---|---|---|
| (Intercept) | 0.030 | [-0.033, 0.092] | 0.349 | 0.031 | 114.669 |
| Mismatch | -0.051 | [-0.106, 0.004] | 0.069 | 0.028 | 10274.426 |
| Strength | -0.028 | [-0.074, 0.018] | 0.232 | 0.023 | 10278.012 |
| Mismatch x Strength | 0.066 | [-0.016, 0.147] | 0.113 | 0.041 | 10266.351 |

**Supplementary Table 9: PC2 scores and RTs**

| Fixed effects | Estimate | CI | p-value | SE | DF |
|---|---|---|---|---|---|

| | | | | | |
|---|---|---|---|---|---|
| (Intercept) | -0.658 | [-0.789, -0.526] | 0.005 | 0.020 | 1.430 |
| PC2 score | 0.007 | [-0.001, 0.014] | 0.077 | 0.004 | 258.877 |
| Strength | 0.208 | [0.193, 0.223] | 0.000 | 0.008 | 108.689 |
| Mismatch | 0.159 | [0.142, 0.176] | 0.000 | 0.009 | 140.013 |
| PC2 score x Strength | 0.008 | [-0.003, 0.018] | 0.163 | 0.005 | 9256.534 |
| PC2 score x Mismatch | 0.004 | [-0.008, 0.017] | 0.504 | 0.006 | 9766.249 |
| Strength x Mismatch | -0.127 | [-0.146, -0.109] | 0.000 | 0.009 | 10204.415 |
| PC2 score x Strength x Mismatch | -0.012 | [-0.031, 0.006] | 0.202 | 0.009 | 9982.026 |

**Supplementary Table 10: PC2 scores as a function of Mismatch, Strength, and Memory Accuracy**

| Fixed effects | Estimate | CI | p-value | SE | DF |
|---|---|---|---|---|---|
| (Intercept) | 0.022 | [-0.038, 0.083] | 0.464 | 0.031 | 126.781 |
| Mismatch | -0.050 | [-0.105, 0.005] | 0.073 | 0.028 | 12075.251 |
| Accuracy | -0.152 | [-0.291, -0.013] | 0.032 | 0.071 | 12122.422 |
| Strength | -0.024 | [-0.070, 0.022] | 0.303 | 0.023 | 12083.804 |
| Mismatch x Accuracy | 0.090 | [-0.119, 0.300] | 0.398 | 0.107 | 12147.439 |
| Mismatch x Strength | 0.061 | [-0.021, 0.142] | 0.144 | 0.042 | 12073.577 |
| Accuracy x Strength | 0.203 | [0.050, 0.355] | 0.009 | 0.078 | 12146.127 |
| Mismatch x Accuracy x Strength | -0.122 | [-0.371, 0.127] | 0.338 | 0.127 | 12115.314 |

**Supplementary Table 11: PC6 scores as a function of Mismatch and Strength**

| Fixed effects | Estimate | CI | p-value | SE | DF |
|---|---|---|---|---|---|
| (Intercept) | 0.020 | [-1.241, 1.281] | 0.878 | 0.103 | 1.015 |
| Mismatch | 0.085 | [0.033, 0.136] | 0.001 | 0.026 | 488.307 |
| Strength | -0.038 | [-0.094, 0.019] | 0.190 | 0.028 | 114.084 |
| Mismatch x Strength | 0.042 | [-0.030, 0.114] | 0.256 | 0.037 | 10240.374 |

**Supplementary Table 12: PC6 scores and RTs**

| Fixed effects | Estimate | CI | p-value | SE | DF |
|---|---|---|---|---|---|
| (Intercept) | -0.653 | [-0.879, -0.428] | 0.014 | 0.026 | 1.198 |
| PC6 score | -0.040 | [-0.052, -0.029] | 0.000 | 0.006 | 145.644 |
| Strength | 0.203 | [0.188, 0.218] | 0.000 | 0.008 | 105.857 |
| Mismatch | 0.158 | [0.141, 0.174] | 0.000 | 0.008 | 139.184 |
| PC6 score x Strength | 0.005 | [-0.006, 0.016] | 0.350 | 0.006 | 4707.178 |
| PC6 score x Mismatch | 0.015 | [0.002, 0.027] | 0.026 | 0.007 | 5396.250 |
| Strength x Mismatch | -0.124 | [-0.142, -0.105] | 0.000 | 0.009 | 10160.143 |
| PC6 score x Strength x Mismatch | -0.008 | [-0.026, 0.011] | 0.415 | 0.009 | 10218.930 |
